# HLTF Prevents G4 Accumulation and Promotes G4-induced Fork Slowing to Maintain Genome Stability

**DOI:** 10.1101/2023.10.27.563641

**Authors:** Gongshi Bai, Theresa Endres, Ulrike Kühbacher, Briana H. Greer, Emma M. Peacock, Magdalena P. Crossley, Ataya Sathirachinda, David Cortez, Brandt F. Eichman, Karlene A. Cimprich

## Abstract

G-quadruplexes (G4s) form throughout the genome and influence important cellular processes, but their deregulation can challenge DNA replication fork progression and threaten genome stability. Here, we demonstrate an unexpected, dual role for the dsDNA translocase HLTF in G4 metabolism. First, we find that HLTF is enriched at G4s in the human genome and suppresses G4 accumulation throughout the cell cycle using its ATPase activity. This function of HLTF affects telomere maintenance by restricting alternative lengthening of telomeres, a process stimulated by G4s. We also show that HLTF and MSH2, a mismatch repair factor that binds G4s, act in independent pathways to suppress G4s and to promote resistance to G4 stabilization. In a second, distinct role, HLTF restrains DNA synthesis upon G4 stabilization by suppressing PrimPol-dependent repriming. Together, the dual functions of HLTF in the G4 response prevent DNA damage and potentially mutagenic replication to safeguard genome stability.

## Introduction

DNA-damaging agents, protein-DNA complexes, and noncanonical nucleic acid secondary structures can threaten genome stability.^1–4^ Among the latter are G-quadruplexes (G4s), nucleic acid structures that form upon the three-dimensional stacking of planar guanidine tetrads.^5,6^ G4s form in G-rich regions of the genome,^5–8^ and are thought to play physiological roles in transcription,^9–11^ telomere homeostasis,^12–15^ DNA replication initiation,^16^ and epigenetic inheritance.^17,18^ Deregulated G4 formation and stabilization, however, can challenge genome stability, by inhibiting DNA replication^19–21^ and transcription^22–24^ ultimately causing DNA damage in the form of DSBs.^25–27^ Indeed, G4-forming motifs are associated with recurrent somatic mutations and are determinants of mutagenesis in the cancer genome.^28^ To mitigate the adverse effects of G4s, cells utilize several proteins to suppress these structures, including helicases that can unwind G4s, as well as ssDNA binding proteins.^6,29^ In addition, the MSH2/MSH6 (MutSα) and MSH2/MSH3 (MutSβ) complexes, whose canonical role is in DNA mismatch recognition and repair,^30^ can also bind to G4s and regulate their stability.^31–33^

When deregulated, G4s pose a barrier to replication fork progression^19–21^ that amongst a variety of other endogenous and exogenous factors can lead to DNA replication stress.^34^ Such obstacles can ultimately cause DNA damage and degradation of the nascent DNA.^35–39^ Cells have elaborate mechanisms to detect and respond to replication stress, including pathways that allow them to tolerate the stress, accomplished by the bypass of replication fork barriers and their subsequent repair or resolution.^39^ Among these DNA damage tolerance (DDT) pathways are replication fork reversal, repriming of DNA synthesis using the alternative primase-polymerase (PrimPol) and translesion synthesis (TLS).^39^ While fork reversal is considered a high-fidelity form of DNA damage tolerance that allows the use of the sister chromatid for lesion bypass, PrimPol-mediated repriming and TLS can be error-prone.^40,41^ How cells choose between these pathways is of great interest.

First identified as a potential transcription factor,^42,43^ helicase-like transcription factor (HLTF), the human ortholog of yeast Rad5, regulates several steps of DDT using all of its known activities.^44^ Using its RING-domain dependent ubiquitin ligase activity, HLTF mediates PCNA ubiquitination to control pathway choice.^45–48^ HLTF can also reverse replication forks. This process is proposed to utilize the ATPase activity of HLTF for translocation on dsDNA and is directed by the HIRAN domain, which binds the 3’-end of ssDNA on the nascent leading strand at a stalled replication fork to position HLTF.^49–53^ Importantly, HLTF-dependent fork reversal leads to the slowing of replication fork progression following replication stress and prevents PrimPol-dependent unrestrained DNA synthesis.^54^ This event is also observed upon loss of other proteins involved in fork reversal.^55^

HLTF may participate in other DNA damage repair pathways as well, promoting efficient nucleotide excision repair (NER) using its translocase activity to evict the incised DNA lesion.^56^ In this process, the HIRAN domain recruits HLTF to the lesion sites before ATPase-dependent eviction of the damaged DNA. Intriguingly, HLTF also has a strand invasion activity dependent on its ATPase domain.^57^ Furthermore, it can displace ssDNA in a triplex structure by translocating on the dsDNA in an ATPase-dependent manner.^49^ How these activities of HLTF relate to its processing of DNA structures *in vivo* is unknown.

Importantly, HLTF has been implicated in tumor suppression and its loss is associated with poor prognosis.^58^ Reduced HLTF expression resulting from promoter hypermethylation is frequently observed in gastric and colorectal cancers and promotes tumor cell growth.^59,60^ Moreover, the knockout of HLTF in mice promotes genome instability and intestinal carcinogenesis.^61^ How HLTF suppresses tumor cell growth and genome instability is not clear.

Here, we uncover an unexpected function for HLTF in maintaining genome stability through G4 regulation. We find that HLTF suppresses the formation of G4s in a MSH2-independent pathway. We find this function of HLTF depends on its ATPase activity and can occur throughout the cell cycle, contrasting with HLTF’s functions during the DNA replication stress response. We also demonstrate that HLTF is enriched at G4s in the human genome and that its binding to chromatin depends largely on transcription. Consistent with its role in preventing G4 accumulation, we find that HLTF prevents alternative lengthening of telomeres (ALT), a process promoted by G4 stabilization. Interestingly, HLTF-deficient cells tolerate the increased replication stress caused by G4 stabilization using PrimPol. Nevertheless, these cells are sensitive to G4 ligands, and this sensitivity is further enhanced in the absence of MSH2. Taken together, our studies reveal a function for HLTF in suppressing the formation of G4s throughout the cell cycle. Furthermore, our studies also show that HLTF loss helps cells to tolerate the formation of these structures during S phase, thereby increasing the opportunities for DNA damage and mutagenesis.

## Results

### HLTF and MSH2 bind chromatin in a reciprocal and cell cycle-independent manner

We previously showed that HLTF restrains replication fork progression in response to DNA replication stress,^52^ and in its absence, DNA synthesis continues in a PrimPol-dependent manner.^54^ To further understand this “stress-resistant” form of DNA synthesis, we examined the proteomic composition of nascent DNA using isolation of proteins on nascent DNA (iPOND) combined with SILAC quantitative mass spectrometry (SILAC-MS).^62,63^ We challenged wild-type (WT) and HLTF-knock-out (HLTF-KO) HEK293T cells with a low dose of hydroxyurea (HU) to induce DNA replication stress and pulse-labeled the nascent DNA with the thymidine analogue EdU (Figure 1A). To isolate equivalent amounts of nascent DNA, we increased the EdU labeling time in WT cells which have slower fork progression than HLTF-KOs (Figure S1A). We then analyzed the enrichment between HLTF-KO and WT samples, identifying proteins with more than a 50% change (Figure 1B). The core components of the replisome were equally enriched on nascent DNA in the two conditions (Figure 1C). By contrast, we observed a more than 50% increase in the enrichment of several proteins involved in DNA mismatch repair (MMR) on HLTF-deficient nascent chromatin, including all components of the MutS complexes (Figure 1B, C).

**Figure 1.**
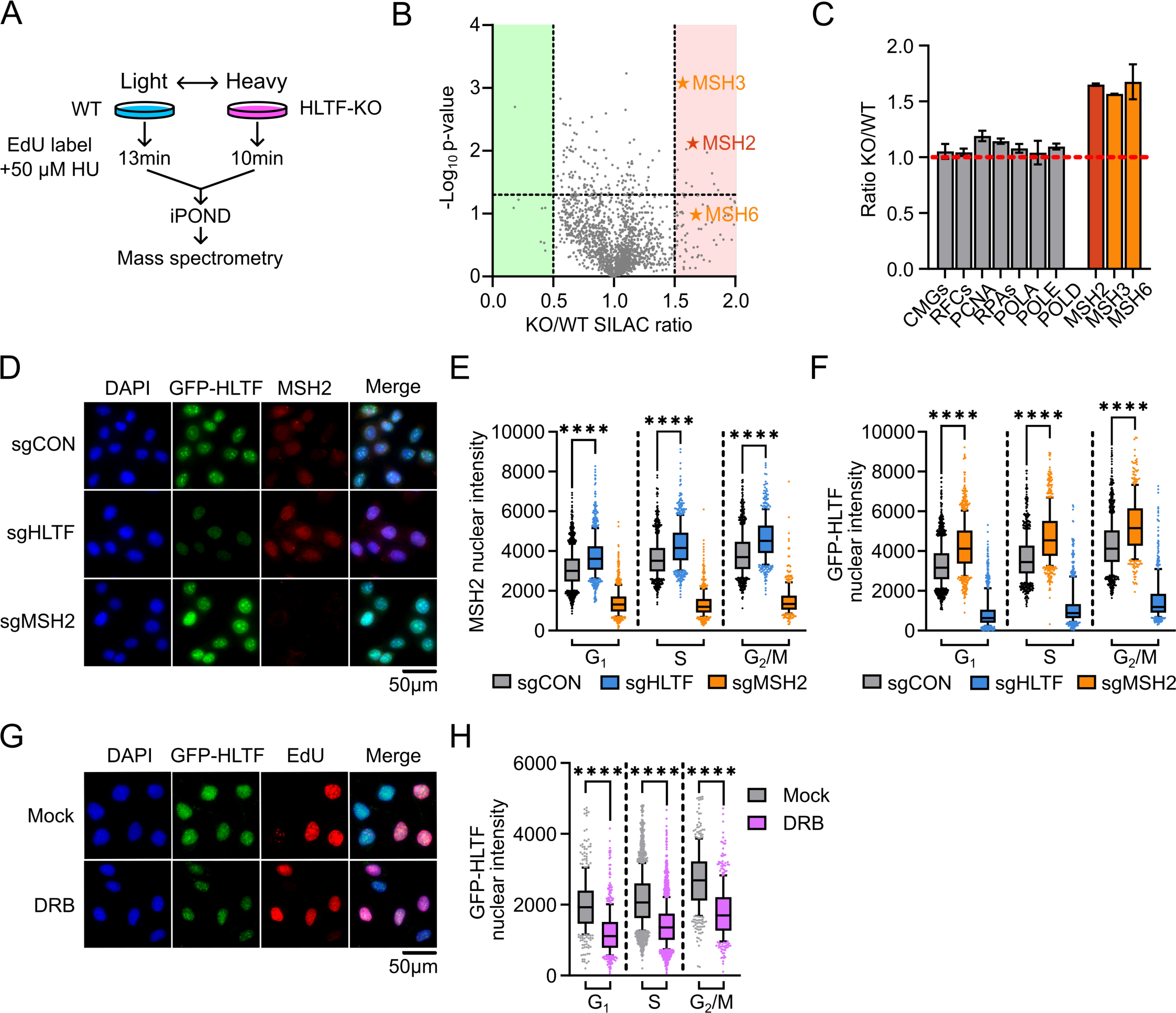
The cell cycle independent regulation of HLTF chromatin binding by MSH2 and transcription. A. Schematic of iPOND-SILAC-MS. “Heavy” and “light” amino acids labeling of WT or HLTF-KO cells were reversed for the two biological repeats. B. Volcano plots of the p-values versus the HLTF-KO/WT SILAC ratio for proteins identified by iPOND-SILAC-MS. p-values are calculated based on two biological repeats. Protein abundance changes with at least 50% decrease (green) or increase (red) are highlighted. Significance cutoff for protein enrichment was set at p=0.05 (dotted line). C. Bar graph showing the HLTF-KO/WT SILAC ratio for iPOND-SILAC-MS identified proteins from B. For CMGs, RFCs, RPAs, and replicative polymerases, mean ± SEM is calculated from the normalized SILAC ratio of each subunit comprising the complex. For MSH2, 3 and 6, the normalized SILAC ratio is used to calculate the mean ± SD (n=2). D. Representative immunofluorescence (IF) images of U2OS cells expressing GFP-HLTF after sgRNA-mediated knockdown. GFP-HLTF is detected using a GFP antibody. E. Box plot showing the mean intensity of chromatin-bound MSH2 protein levels as shown in D. ∗∗∗∗ p<0.0001, by Mann-Whitney test. See also Figure S1E. F. Box plot showing the mean intensity of chromatin-bound GFP-HLTF protein levels as shown in D. ∗∗∗∗ p<0.0001, by Mann-Whitney test. See also Figure S1G. G. Representative IF images of U2OS cells expressing GFP-HLTF after 4h of mock or DRB treatment (100 µM). GFP-HLTF is detected using a GFP antibody. H. Box plot showing the mean intensity of chromatin-bound GFP-HLTF protein levels after 4h of mock or DRB treatment (100 µM) as shown in G. ∗∗∗∗ p<0.0001, by Mann-Whitney test. See also Figure S1K. For all IF stainings, cells are pre-extracted to retain the chromatin-bound protein. For all box plots, whiskers indicate the 10th and 90th percentiles, boxes span the 25th to 75th percentiles. Lines inside boxes represent medians.

MSH2 interacts with MSH3 or MSH6 to form the MutSα or β complexes, respectively. These proteins are known to bind the nascent DNA during S/G_2_ repairing mismatches formed during DNA synthesis.^30^ Hence, we hypothesized that they might be acting on mismatches that result from the action of PrimPol and other TLS polymerases in HLTF’s absence.^54^ To test this idea and validate our iPOND data, we used quantitative image-based cytometry (QIBC) in pre-extracted U2OS cells to analyze the association of MSH2 with chromatin expressing endogenously tagged GFP-HLTF (Figure S1B).^56^ We pulse-labeled cells with EdU to identify cells in different cell cycle phases based on their total and nascent DNA content. We observed increased association of MSH2 with chromatin upon knocking down HLTF (Figure 1D; Figure S1C, D). Unexpectedly, this increase was observed not only in S and G_2_/M phase cells but also in G_1_ cells (Figure 1E; Figure S1E). This suggests that HLTF suppresses the binding of MSH2 to chromatin throughout the cell cycle and is inconsistent with the idea that these proteins are responding to mismatches introduced post-replicatively.

Next, we asked if the loss of MSH2 alters the association of HLTF with chromatin. Surprisingly, when we knocked down MSH2 in the GFP-HLTF cell line, we detected increased HLTF chromatin binding (Figure 1D, F; Figure S1C, F, G). The knockdown of MSH2 in wildtype U2OS cells also led to increased HLTF chromatin binding (Figure S1H, I). Intriguingly, we observed increased chromatin binding of HLTF in all cell cycle phases (Figure 1F; Figure S1H, I). These results suggest that HLTF and MSH2 can suppress the interaction of the other with chromatin throughout the cell cycle.

The binding of HLTF to chromatin throughout the cell cycle and the previous links of HLTF to transcription,^42,43^ prompted us to ask if its chromatin binding is dependent on transcription. We found that the inhibition of transcription with 5,6-dichlorobenzimidazole (DRB) reduced HLTF chromatin binding in all cell cycle phases (Figure 1G, H; Figure S1J, K). These results indicate that the interaction of HLTF with chromatin can occur in a transcription-dependent and replication-independent manner that is increased when MSH2 is lost. They also suggest that HLTF and MSH2 interact with chromatin in a reciprocal manner.

### HLTF prevents the accumulation of G4 structures

We next wanted to determine the function of HLTF and MSH2 on chromatin. Nucleic acid secondary structures such as G4s and RNA-DNA hybrids can form co-transcriptionally.^64–66^ In addition to binding DNA mismatches,^30^ MutSα and β complexes can interact with some nucleic acid secondary structures,^67–71^ including G4s.^31–33^ Given the interplay between HLTF and MSH2 and the reduction in HLTF’s chromatin binding upon transcription inhibition, we asked if HLTF controls G4 levels *in vivo.* We observed an increase in G4 signal in HLTF-KO cell lines using two different cell types and two distinct antibodies raised against G4-structured DNA (Figure 2A, B; Figure S2A-D). We also found that transient knockdown of HLTF in U2OS cells increased the G4 signal (Figure S2E). Interestingly, this increase was observed in all cell cycle phases, consistent with the effect of HLTF loss on MSH2 chromatin binding.

**Figure 2.**
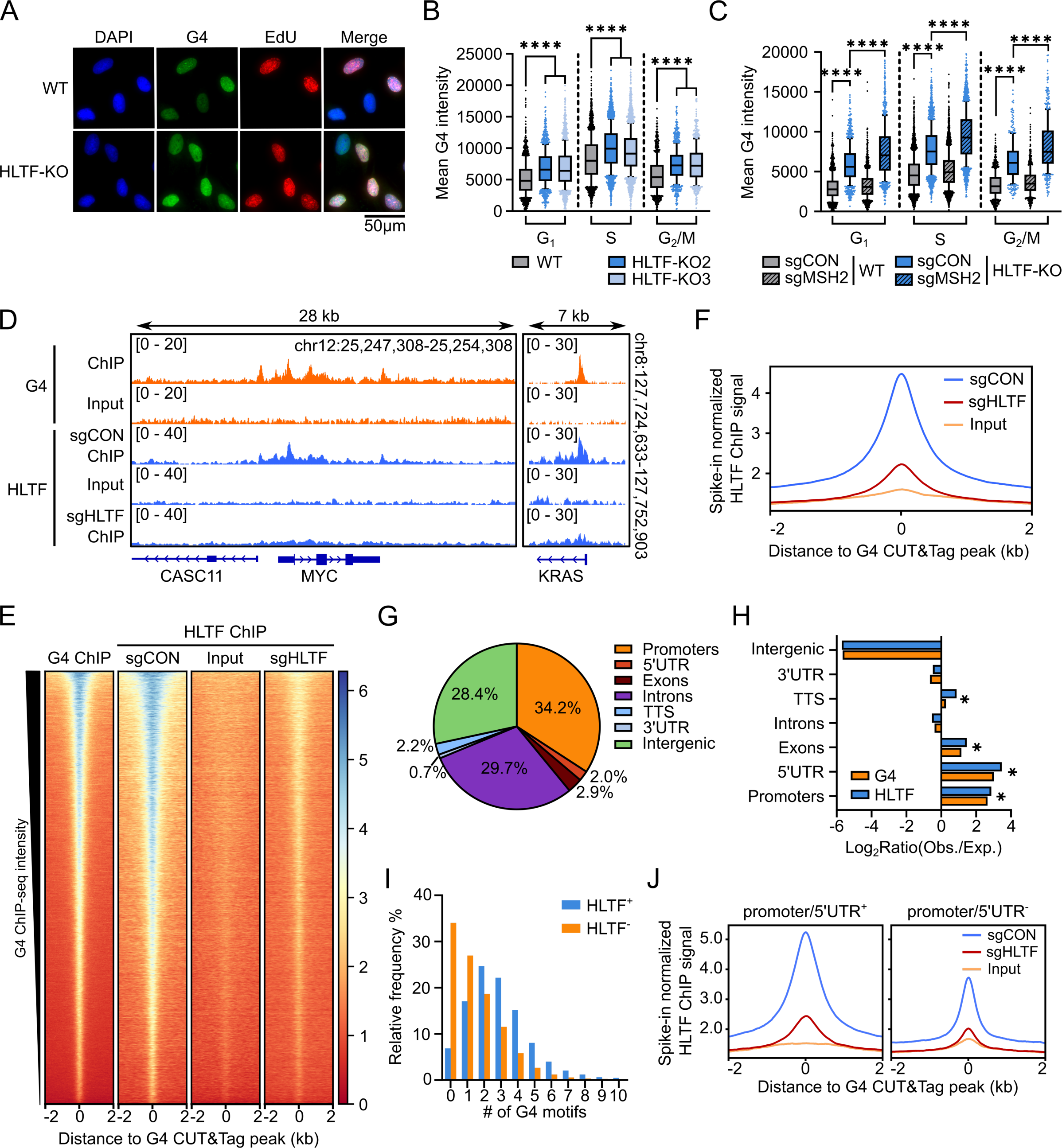
HLTF suppresses G4 accumulation in the cells and is enriched at G4 structures in the human genome. A. Representative G4 IF images in WT and isogenic HLTF-KO U2OS cells. G4s are detected using the 1H6 antibody. B. Box plot showing the mean G4 intensity in WT and isogenic HLTF-KO U2OS cells, as shown in A. ∗∗∗∗ p<0.0001, by Mann-Whitney test. See also Figure S2C. C. Box plot showing the mean G4 intensity in WT and isogenic HLTF-KO U2OS cells after sgRNA-mediated knockdown. ∗∗∗∗ p<0.0001, by Mann-Whitney test. See also Figure S2F. For all box plots, whiskers indicate the 10th and 90th percentiles, boxes span the 25th to 75th percentiles. Lines inside boxes represent medians. D. Representative browser tracks of the *MYC* and *KRAS* loci from a G4 ChIP-seq in U2OS cells (GSE162299) and HLTF ChIP-seq in U2OS cells expressing GFP-HLTF. E. Heatmaps showing ChIP-seq coverage for G4 and HLTF at G4 CUT&Tag peaks (n=35,104)(GSE181373). The x-axis represents the distance from the G4 CUT&Tag peak in kb. Heatmaps were sorted by G4 ChIP-seq signal intensity for all ChIP-seq samples. Spearman correlation coefficient between G4 and HLTF sgCON ChIP-seq signal was 0.74. F. Aggregate plot showing HLTF ChIP-seq coverage (y-axis) relative to the distance (x-axis) from G4 CUT&Tag peaks in U2OS cells in kb, related to E. G. Pie chart showing the distribution of HLTF ChIP-seq peaks within each genomic compartment in the human genome. TTS, transcription termination site. H. Bar graph showing the relative enrichment of G4 CUT&Tag and HLTF ChIP-seq peaks within each genomic compartment in the human genome. ∗ indicates compartments where both G4 and HLTF showed significant enrichment. I. Histogram showing the frequency distribution of the number of G4 motifs within G4 CUT&Tag peaks. G4 peaks are segregated based on if they overlap with a HLTF ChIP-seq peaks: G4 peaks that overlap with HLTF peaks are HLTF^+^ or otherwise HLTF^-^. See also Figure S2H. J. Aggregate plot showing HLTF ChIP-seq coverage (y-axis) relative to the distance (x-axis) from G4 CUT&Tag peaks in U2OS cells in kb. G4 peaks are segregated based on whether they are within the promoter or 5’UTR (G4 in promoter/5’UTR or promoter/5’UTR^+^, n=17,620; G4 out of promoter/5’UTR or promoter/5’UTR^-^, n=17,484). Promoters are identified as ±1kb of the annotated transcription start site. See also Figure S2I.

Given the reciprocal relationship between HLTF and MSH2 chromatin binding, we also asked whether knocking down MSH2 increased the G4 signal and whether the proteins might interact at a functional level in the same or different pathways. We observed a modest increase in the G4 signal when MSH2 alone was knocked down. However, we observed a synergistic increase in the G4 signal when MSH2 was knocked down in an HLTF-KO U2OS cell line (Figure 2C; Figure S2F). These effects were observed in all cell cycle phases. Taken together, our data suggest that HLTF and MSH2 may act in independent pathways to suppress the accumulation of G4s throughout the cell cycle.

### HLTF binds to G4-containing loci throughout the genome

The ability of HLTF to suppress G4 accumulation in cells raised the possibility that it might act at G4s in the genome. To test this hypothesis, we analyzed HLTF binding on chromatin at sites known to form G4s. To accurately evaluate changes in HLTFs chromatin association, we performed spike-in normalized chromatin immunoprecipitation-sequencing (ChIP-seq)^72^ in U2OS cells with endogenously tagged GFP-HLTF using an antibody against GFP. We included a HLTF knockdown to ensure the specificity of the ChIP-seq signal.

First, we examined the enrichment of HLTF ChIP-seq signal over input and HLTF knockdown at G4-forming sites (Figure 2D, E) previously identified in U2OS cells using two different methods (ChIP-seq and CUT&Tag).^11,73^ Ranking HLTFs chromatin binding based on the G4 ChIP-seq signal intensity at G4s identified by CUT&Tag, we found that HLTF binds to G4 sites throughout the genome and that its enrichment correlates with the G4 signal at these sites (Figure 2E, F).

To further assess the ability of HLTF to bind *bona fide* G4 structures, and not simply the underlying G-rich sequences, we analyzed HLTF enrichment at G4 forming motifs that have been observed to form G4s *in vitro*.^74^ Importantly, only a subset of these motifs form G4s in cells, when identified by G4 CUT&Tag.^73^ We found that HLTF is specifically enriched at those motifs that actually form G4s in cells and not at those that only form *in vitro* (Figure S2G). Hence, HLTF is enriched at *bona fide* G4 structures in cells.

Next, we identified HLTF binding sites in the genome, calling 7,614 HLTF-specific ChIP peaks. HLTF peaks are predominantly found at the beginning of genes, including promoters and 5’UTRs (Figure 2G, H). Interestingly, HLTF’s enrichment in the genome is similar to that observed for G4s (Figure 2H).^9^ Moreover, we found that 56% (n=4,261) of HLTF peaks overlap with G4-forming sites (Figure S2H) and were also more likely to contain multiple G4-forming motifs (Figure 2I).^74^ Given the enrichment of HLTF in promoters and 5’UTRs, we asked if its enrichment at G4s is simply due to its binding to transcriptional regulatory regions. We found, however, that HLTF binds to G4s that form outside of a promoter or 5’UTR. Although this enrichment was lower, it still correlates with the G4 intensity (Figure 2J; Figure S2I). Taken together, these data demonstrate that HLTF is enriched at G4 structures formed throughout the genome and its binding is a function of G4 density, consistent with its role in regulating G4 levels in cells.

### HLTF is enriched at G4s stabilized by an R-loop

G4 DNA structures can co-occur with R-loops at transcriptionally active regions and at telomeres,^15,75^ with each structure stabilizing formation of the other.^64,75,76^ Hence, we asked if HLTF is found at sites of RNA-DNA hybrid formation. We examined the enrichment of HLTF ChIP-seq signal at RNA-DNA hybrid sites previously identified by DRIP-seq in U2OS cells.^26^ Interestingly, we found only a modest enrichment of HLTF at RNA-DNA hybrid forming sites relative to that observed at G4s (Figure 3A). However, when we examined the 15% of RNA-DNA hybrid sites that overlap with G4s identified by G4 CUT&Tag, we saw substantial enrichment of HLTF relative to its enrichment at hybrids without a G4 (Figure 3A, B; Figure S3A). We also found that HLTF’s enrichment at G4 sites is stronger when the G4 site overlaps with RNA-DNA hybrids (Figure S3B).

**Figure 3.**
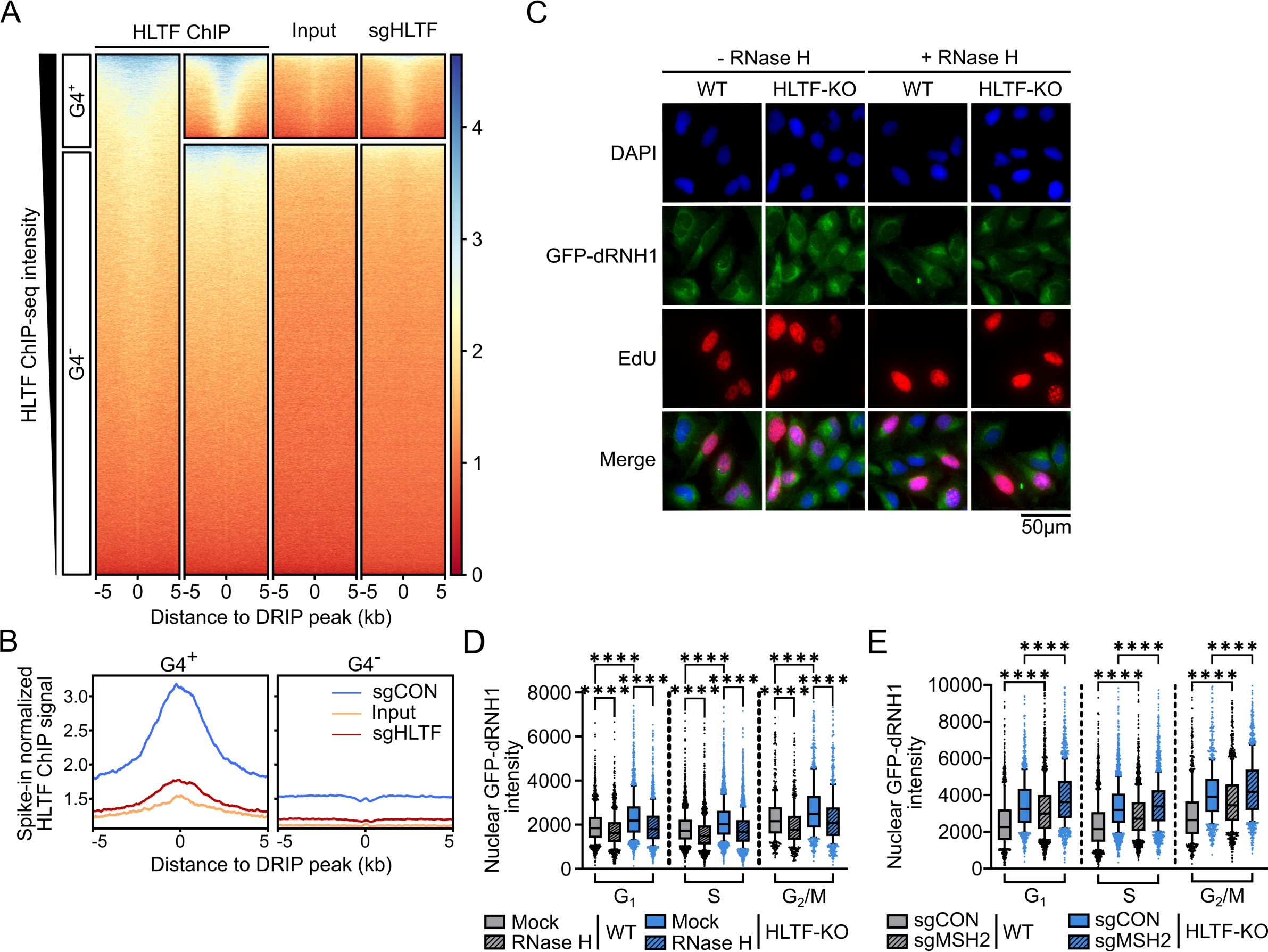
HLTF is enriched at G4s stabilized by RNA-DNA hybrids. A. Heatmaps showing ChIP-seq coverage for HLTF at RNA-DNA hybrid peaks identified in U2OS cells by DRIP-seq (GSE115957). The x-axis represents the distance from the DRIP peak in kb. DRIP peaks are segregated based on whether they overlap with G4 peaks identified by G4 CUT&Tag: DRIP peaks that overlap with G4 peaks are G4^+^(n=11,499) or otherwise G4^-^ (n=59,090). B. Aggregate plot showing HLTF ChIP-seq coverage (y-axis) relative to the distance in kb (x-axis) from DRIP peaks identified in U2OS cells. Related to A. See also Figure S3A. C. Representative IF images of RNA-DNA hybrids in WT and isogenic HLTF-KO U2OS cells after mock or RNase H digestion. RNA-DNA hybrids are then detected using the GFP-dRNH1 reagent. D. Box plot showing the mean nuclear RNA-DNA hybrid intensity, as shown in C. ∗∗∗∗ p<0.0001, by Mann-Whitney test. See also Figure S3C. E. Box plot showing the mean nuclear RNA-DNA hybrid intensity in WT and isogenic HLTF-KO U2OS after sgRNA transfection. ∗∗∗∗ p<0.0001, by Mann-Whitney test. See also Figure S3D. For all box plots, whiskers indicate the 10th and 90th percentiles, boxes span the 25th to 75th percentiles. Lines inside boxes represent medians.

To test the relationship between HLTF and RNA-DNA hybrids further, we measured the accumulation of RNA-DNA hybrids in cells using GFP-tagged catalytically-inactive RNase H1 as a specific probe for RNA-DNA hybrids.^77^ We observed a small but significant RNaseH-reversible increase in hybrid signal in HLTF-KO cells (Figure 3C, D; Figure S3C). Knockdown of MSH2 alone also modestly increased the RNA-DNA hybrid signal, and as observed for G4s, the knockdown of MSH2 in HLTF-KO cells led to a further increase in all cell cycle phases (Figure 3E; Figure S3D). These observations are consistent with our finding that HLTF binds the subset of RNA-DNA hybrid sites that also coincide with G4s on chromatin (Figure 3A). Taken together, our data indicates that HLTF’s binding to chromatin is dependent on its association with G4s, and that HLTF only binds and acts on those R-loops that also contain and are stabilized by a G4.

### HLTF suppresses G4 formation in an ATPase-dependent manner

Next, we sought to determine the mechanism by which HLTF prevents the accumulation of G4s. HLTF contains three known domains: the HIRAN domain, a RING domain containing ubiquitin ligase, and an ATPase domain (Figure 4A). We previously showed that mutation of HLTF’s HIRAN domain (R71E) prevents 3’-OH DNA end binding and fork remodeling, but does not alter its ATPase and ubiquitin ligase activities.^52,54^ To investigate the role of each of these domains in G4 suppression, we generated mutant forms of HLTF predicted to lack these activities. We generated the C760S mutant by modifying a residue in the RING domain required for coordinated zinc binding and ubiquitin ligation,^78^ and the R890Q mutant by modifying a residue analogous to R764 in the related replication fork remodeler, SMARCAL1 (Figure S4A). The analogous mutation is observed in Schimke immuno-osseous dysplasia disease patients and disrupts the ATPase activity of SMARCAL1.^79^

**Figure 4.**
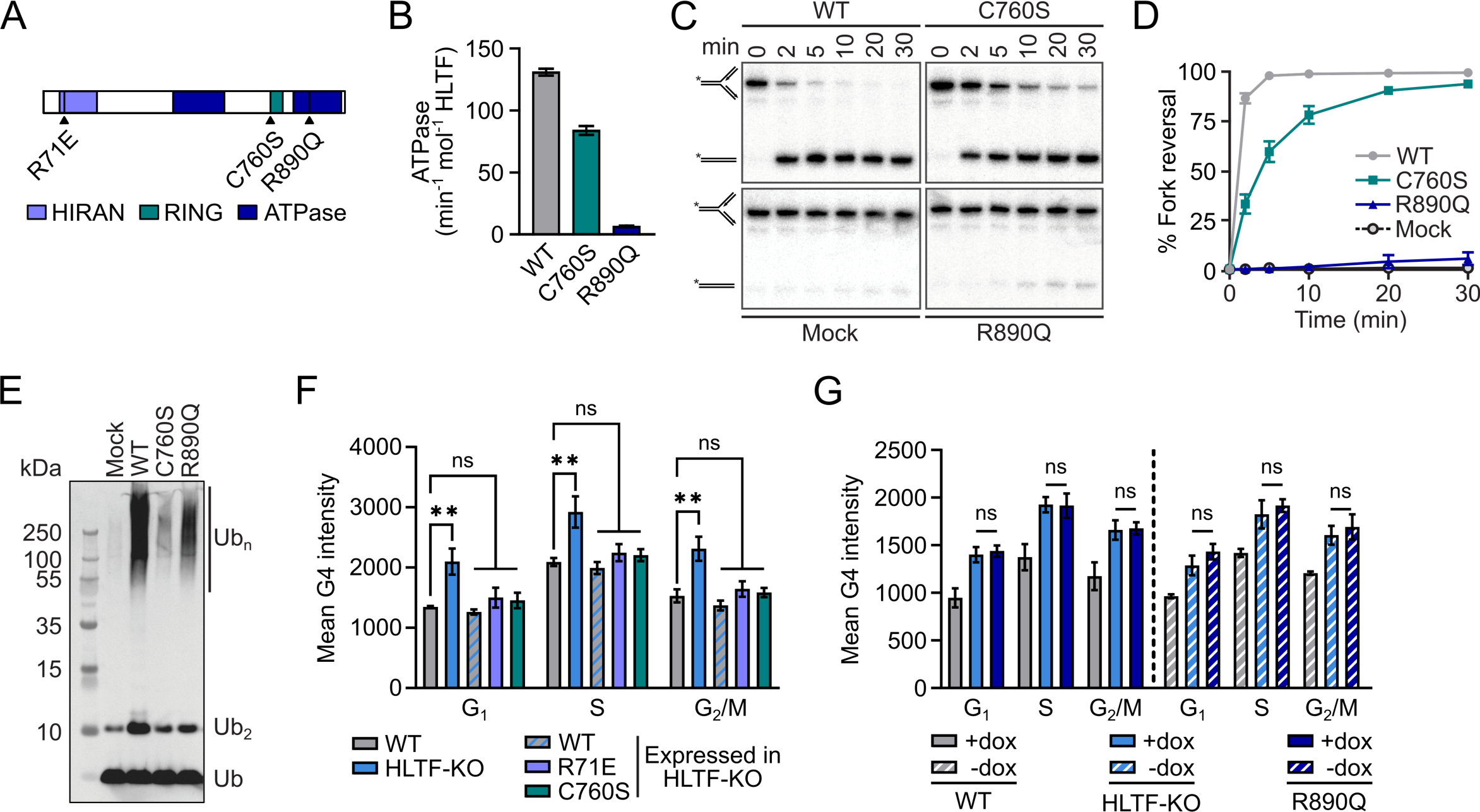
HLTF suppresses G4s in an ATPase-dependent manner in cells. A. Schematic of HLTF domain structures. Lines inside each domain represent the position of the mutations. B. Bar graph showing the ATPase rates of HLTF WT and mutant proteins measured using an NADH-coupled assay. Rates were corrected for background NADH decomposition in a no-enzyme control. Data are plotted as the mean ± SD (n=3). See also Figure S4B. C. Denaturing PAGE showing the separation of DNA substrate (fork) and product (duplex) over time, as a measurement of the *in vitro* fork reversal activity of HLTF WT and mutant proteins. D. Dot and line graph showing the *in vitro* fork reversal activity of HLTF WT and mutant proteins as a function of time. Related to C. Data are plotted as the mean ± SD (n=3). E. Western blot analysis of HLTF-dependent Ub chain formation by UBC13/MMS2. A ubiquitin antibody is used to detect ubiquitin chain formation. F. Bar graph showing the mean G4 intensity in WT, isogenic HLTF-KO, and HLTF-KO U2OS cells expressing HLTF WT or mutant proteins. Two clones of each HLTF WT or mutant expressing cell lines were analyzed separately. The median of the mean G4 intensity of the two clones is averaged to calculate the mean ± SEM (n=3). ∗∗ p < 0.01; ns, not significant, by one-way ANOVA followed by Dunnett’s test. G. Bar graph showing the mean G4 intensity in WT, isogenic HLTF-KO, and HLTF-KO U2OS cells expressing HLTF R890Q mutant, after dox induction (24 h, 500 ng/mL). Two clones of R890Q expressing HLTF-KO cell lines are analyzed separately. The median of the mean G4 intensity of the two clones is averaged to calculate the mean ± SEM (n=3). ns, not significant, by t-test.

We then tested the biochemical activity of these new mutants to validate their functions and confirm the separation of functions. We found that the C760S mutant retains over 60% of its ATPase activity (Figure 4B; Figure S4B), consistent with its ability to reverse forks *in vitro* (Figure 4C, D). Furthermore, we found that the R890Q mutant lacks both ATPase and fork reversal activities (Figure 4B, C, D; Figure S4B). The ubiquitin ligase activity is lost in the C760S mutant but is retained in the R890Q mutant (Figure 4E).

To determine if the suppression of G4s is dependent on any of these domains, we examined the G4 signal in HLTF-KO cells expressing WT or mutant forms of HLTF. We had previously generated HLTF-KOs stably expressing WT or HIRAN mutant (R71E) proteins,^54^ and we generated cells expressing the C760S mutant using the same system (Figure S4C). We also generated HLTF-KOs inducibly expressing the R890Q mutant protein (Figure S4D).

As expected, the expression of WT HLTF in HLTF-KO cells reduced G4 levels to those found in the parental U2OS cell line. Expression of either the R71E or C760S mutant also suppressed G4 accumulation in HLTF-KO cells in all cell cycle phases (Figure 4F). By contrast, the ATPase mutant, while still able to bind chromatin (Figure S4E), failed to suppress G4s when inducibly expressed in HLTF-KO cells (Figure 4G). These data indicate that HLTF-dependent suppression of G4 accumulation specifically requires its ATPase domain. Thus, HLTF’s ability to prevent G4 accumulation is unlikely related to its effects on replication fork reversal or PCNA ubiquitination, which require the HIRAN^51,52^ and ubiquitin ligase^45,48^ functions, respectively.

### HLTF suppresses ALT activity in an ATPase-dependent manner

G4s and RNA-DNA hybrids both form at telomeres^65,66,80–83^ and their interdependent stabilization has been associated with increased alternative lengthening of telomeres (ALT) activity.^15^ MutS proteins have been shown to suppress ALT activity.^33,84^ The increased accumulation of G4s and RNA-DNA hybrids upon loss of HLTF and MSH2 (Figure 2C; Figure 3E) thus raised the possibility that these proteins may suppress ALT activity.^33,84^

Large RPA foci can be observed at telomeres undergoing ALT.^85^ We observed the increased formation of large RPA foci in ALT-positive U2OS cells after loss of either HLTF or MSH2, and loss of both factors led to a synergistic increase in foci formation (Figure S5A, B). Treatment with the G4-stabilizing ligand pyridostatin (PDS) further increased RPA foci formation in all of these conditions (Figure S5A, B). By contrast, ALT-negative RPE1 cells did not exhibit a significant increase in the formation of RPA foci after similar treatments (Figure S5C, D). These data are consistent with the idea that HLTF and MSH2 have a role at telomeres in ALT-positive cells.

Next, we used telomere FISH and found that ∼90% of the RPA foci colocalized with telomeres in WT and HLTF-KO cells, with more foci forming in HLTF-KOs (Figure S5E, F, G). We then examined several markers of ALT. Telomeres associate with promyelocytic leukemia (PML) bodies during ALT to form ALT-associated PML bodies (APBs).^86^ We observed increased formation of APBs in HLTF-KO cells compared to the parental U2OS cells (Figure 5A, B). Finally, we monitored *in situ* ALT-specific single-stranded telomeric C-rich DNA (ssTelo-C), an independent marker for ALT.^87^ We found that ssTelo-C formation also increased in HLTF-KO cells (Figure 5C, D). These data demonstrate that HLTF suppresses ALT activity.

**Figure 5.**
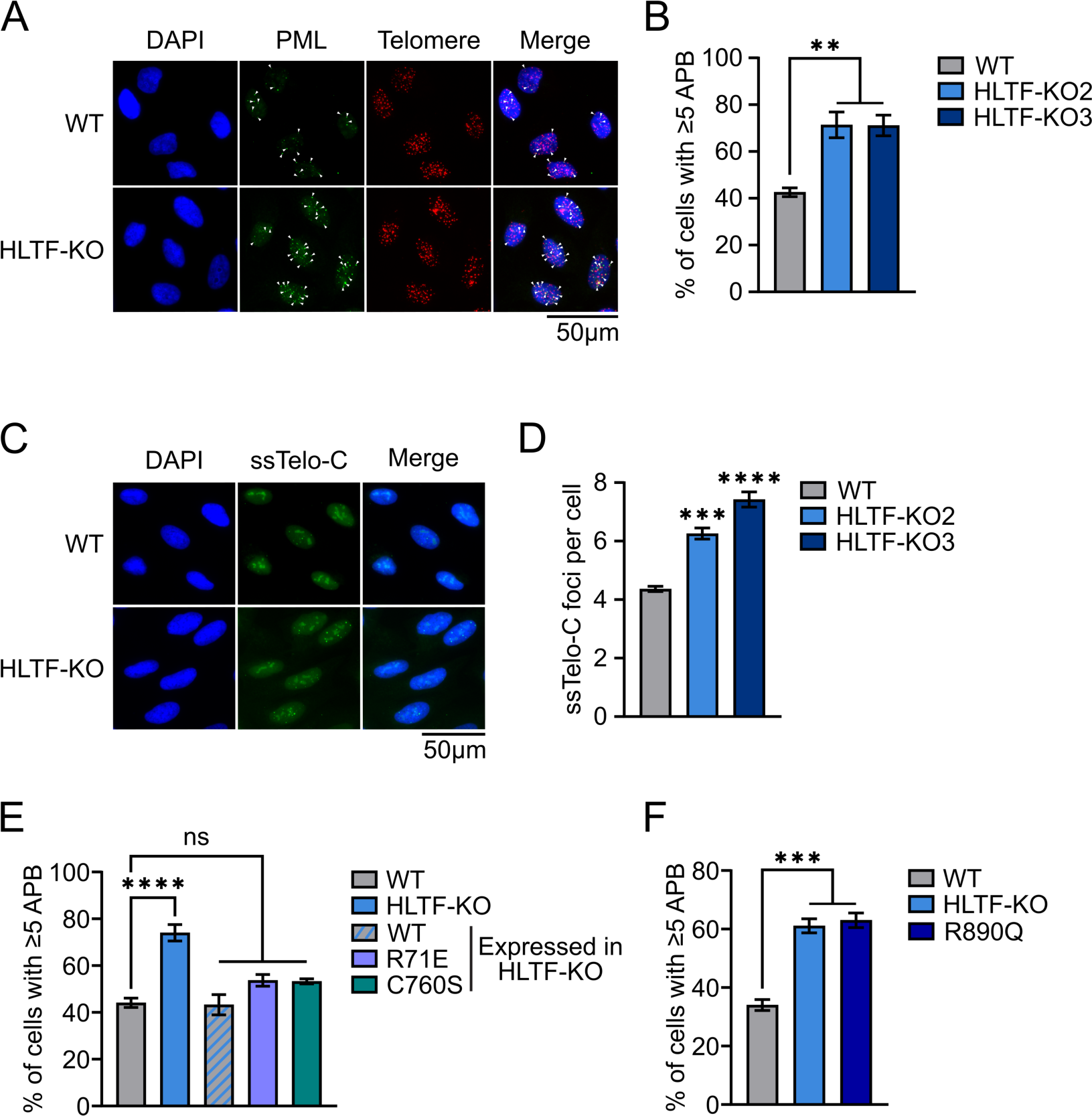
HLTF suppresses ALT activity in an ATPase-dependent manner. A. Representative IF images of ALT-associated PML body (APB) detection in WT and isogenic HLTF-KO U2OS cells. Arrowheads mark the APBs. B. Bar graph showing the percentage of cells with at least five APBs. Related to A. Data are represented as mean ± SEM (n = 3). ∗∗ p < 0.01, by one-way ANOVA followed by Dunnett’s test. C. Representative IF images of *in situ* ssTelo-C foci detection in WT and isogenic HLTF-KO U2OS cells. D. Bar graph showing the average number of ssTelo-C foci per cell. Related to C. Data are represented as mean ± SEM (n = 3). ∗∗∗ p < 0.001; ∗∗∗∗ p < 0.001, by one-way ANOVA followed by Dunnett’s test. Results comparing to the WT samples are shown. E. Bar graph showing the percentage of cells with at least 5 APBs in WT and isogenic HLTF-KO U2OS cells, and HLTF-KO cells expressing HLTF WT, R71E or C760S mutant proteins. Data are represented as mean ± SEM (n ≥ 3). ∗∗∗∗ p < 0.0001; ns, not significant, by one-way ANOVA followed by Dunnett’s test. F. Bar graph showing the percentage of cells with at least 5 APBs after dox induction (500 ng/mL, 24 h) in WT and isogenic HLTF-KO U2OS cells, and HLTF-KO cells expressing the R890Q mutant. Data are represented as mean ± SEM (n = 3). ∗∗∗ p < 0.001, by one-way ANOVA followed by Dunnett’s test.

HLTF’s effect on ALT could be related to its role in controlling fork progression or to its role in preventing the accumulation of G4s and RNA-DNA hybrids. To address this question, we evaluated APB formation in our HLTF mutant cell lines, taking advantage of the different roles for the HIRAN, ubiquitin ligase and ATPase domains. We found that the expression of WT, R71E, or C760S mutants suppressed the formation of APBs (Figure 5E), whereas the expression of the R890Q mutant did not (Figure 5F). Altogether, these data indicate that HLTF reduces ALT activity by suppressing the formation of G4s at telomeres.

### HLTF restrains replication fork progression in response to G4 stabilization

G4 stabilization can inhibit replication fork progression,^19–21,88,89^ and we previously demonstrated that HLTF promotes fork reversal and restrains fork progression in response to replication stress-inducing reagents.^54^ Intriguingly, HLTF-KO cells have increased G4s and RNA-DNA hybrids in the S phase, yet they exhibit fork progression rates similar to the parental U2OS cell line.^52,54^ We thus wondered if loss of HLTF allows cells to tolerate increased levels of these secondary structures. To address this question, we challenged cells with the G4-stabilizing ligand pyridostatin (PDS) and carried out a DNA fiber assay (Figure 6A). We found that PDS treatment inhibited replication fork progression in the parental U2OS cell line. In HLTF-KO cells, however, DNA synthesis continued following PDS treatment at rates similar to untreated cells (Figure 6B). This suggests that HLTF restrains fork progression upon G4 stabilization.

**Figure 6.**
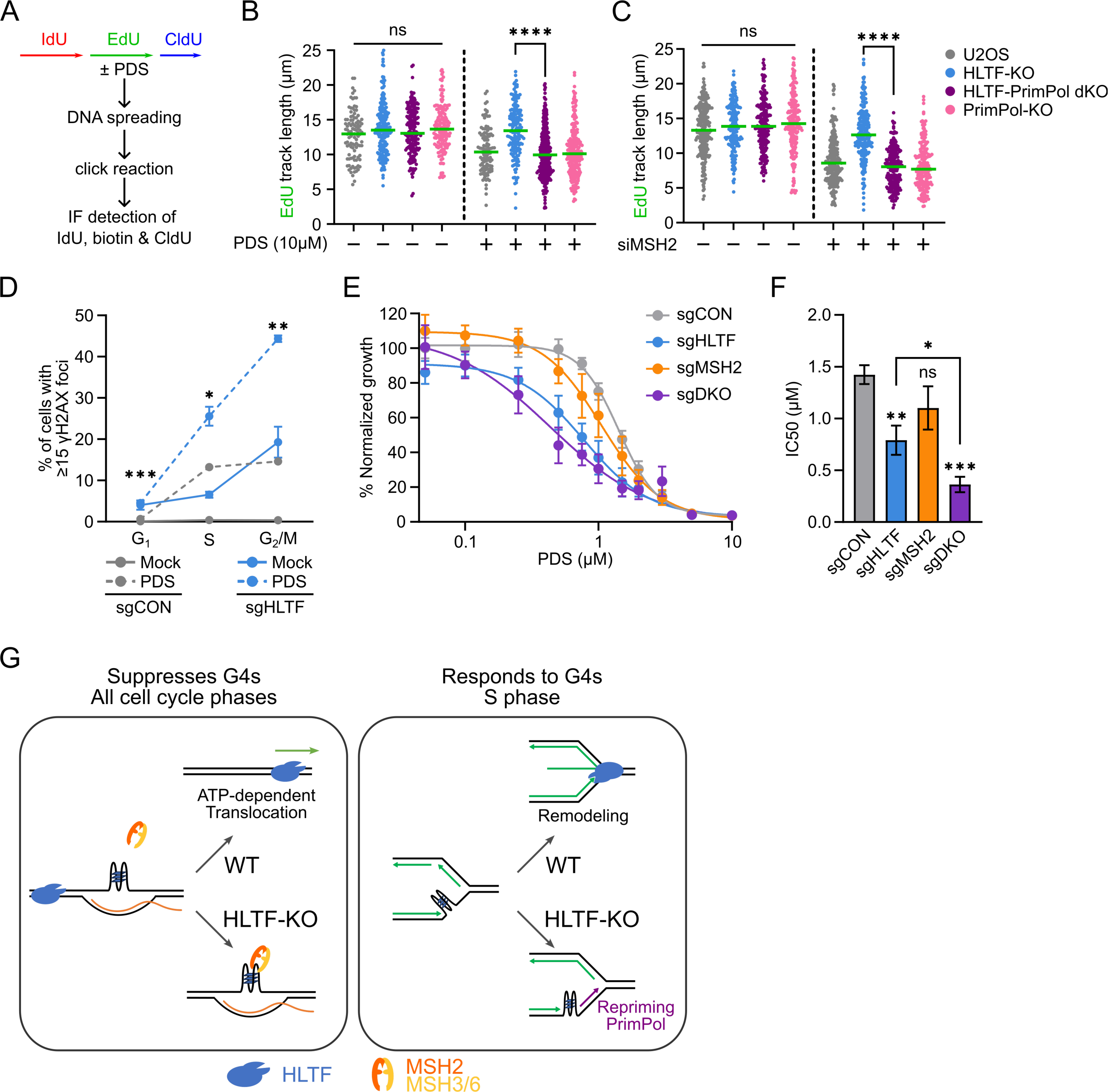
HLTF restrains replication fork progression, protects cells from DNA damage and growth defects in response to G4-stabilization. A. Experimental setup for replication fork progression assay with triple nucleotide labeling. B. Dot plot of EdU tract lengths (n=3) in U2OS cells either mock or treated with PDS. Line represents the median. ns, not significant, by Kruskal-Wallis test; ∗∗∗∗ p < 0.0001, by Mann-Whitney test. C. Dot plot of EdU tract lengths (n=3) in U2OS cells 72h after siRNA transfection. Line represents the median. ns, not significant, by Kruskal-Wallis test; ∗∗∗∗ p < 0.0001, by Mann-Whitney test. D. Dot and line graph showing the percentage of RPE1 cells with at least 15 γH2AX foci from each cell cycle phase after mock or PDS (5 μM) treatment for 24 h after sgRNA transfection. Data are represented as mean ± SEM (n=3). ∗ p < 0.05; ∗∗ p < 0.01; ∗∗∗ p < 0.001, by t-test. Test results comparing the sgCON and sgHLTF PDS-treated samples are shown. E. Drug-response curve for PDS in RPE1 cells after sgRNA transfection, using confluence as a readout after six days of treatment. Cells are transfected with sgRNAs against both HLTF and MSH2 in the sgDKO sample. Data are represented as mean ± SEM (n = 4). F. Bar graph showing the IC50 value of RPE1 cells in response to PDS treatment after sgRNA transfection. Related to E. Data are represented as mean ± SEM (n=3). ∗∗ p < 0.01; ∗∗∗ p < 0.001; ns, not significant, by one-way ANOVA followed by Dunnett’s test compared to sgCON; ∗ p < 0.05 by t-test. G. Proposed model for the dual-roles of HLTF in G4 regulation to maintain genome stability. In its first role, HLTF acts in all cell cycle phases and uses its ATPase-dependent dsDNA translocase activity to travel on dsDNA. Once encountering nucleic acid secondary structure such as a G4, the translocation action of HLTF may directly destabilize the structure while reannealing the structure forming sequence with its complementary strand. Alternatively, HLTF’s translocase activity may simply help localize it to G4s, where it may dismantle the structure using other unknown mechanisms, or recruits other G4 resolution factors to resolve the G4, therefore restoring the G4 structure into duplex DNA. MutS complexes can also bind and regulate G4s in a distinct pathway that is independent of HLTF. In a second role, HLTF restrains DNA synthesis in response to G4 stabilization by promoting fork remodeling. In HLTF’s absence, G4s are bypassed by PrimPol-mediated repriming during DNA replication.

Following HU-induced replication stress, DNA synthesis is sustained in HLTF-deficient cells by PrimPol-mediated repriming.^54^ PrimPol can also bypass G4s formed at replication forks to prevent extensive uncoupling of the replicative helicase and polymerase.^90^ Therefore, we asked whether PrimPol promotes fork progression following PDS treatment when HLTF is absent. Indeed, replication fork progression was reduced dramatically following PDS challenge in PrimPol-HLTF double-knockout (dKO) cells (Figure 6B). These results suggest that PrimPol promotes the bypass of PDS-stabilized G4s at replication forks in the absence of HLTF.

The knockdown of MSH2 elevates the level of G4s and RNA-DNA hybrids in a manner that is enhanced when HLTF is also lost, and both structures are known to slow replication fork progression and cause replication stress.^19,20,81,82,88,91^ Indeed, we found that knocking down MSH2 reduced replication fork progression even in unchallenged conditions (Figure 6C). However, knocking down MSH2 in HLTF-KO cells did not slow fork progression. Furthermore, DNA synthesis was reduced in HLTF-PrimPol dKO cells, indicating it is dependent on PrimPol (Figure 6C). Taken together, these data suggest that HLTF restrains fork progression upon increased formation of secondary DNA structures and that its loss promotes continued, PrimPol-dependent replication fork progression in this context. Hence, in the absence of HLTF, cells can continue DNA synthesis, effectively tolerating a variety of impediments to replication fork progression.

### HLTF protects cells from DNA damage and growth defects induced by G4-ligands

The ability of HLTF-deficient cells to replicate their DNA upon PDS challenge or in the absence of MSH2 led us to ask about the long-term effect of elevated G4 levels on cells. The bypass of G4s with PrimPol might allow continued DNA synthesis but it could leave unresolved G4s, which could prevent the completion of DNA replication and cause DNA damage if not promptly resolved. Yet, we previously found that HLTF-deficient cells exhibit resistance to HU-induced replication stress and accumulate less DNA damage than WT cells under these conditions.^54^

To determine how G4 formation and stabilization affect genome stability and growth of HLTF-deficient cells, we examined the formation of γH2AX foci, a marker for DNA damage. To minimize potentially confounding effects of telomeric damage, we assessed γH2AX foci in ALT-negative p53-deficient RPE1 cells (Figure S5D). The knockdown of HLTF increased G4 formation in these cells in both G_1_ and G_2_/M phase, although the knockdown of MSH2 had little effect on G4s in these cells (Figure S6A). Interestingly, upon knockdown of HLTF, we also observed an increase in the formation of γH2AX foci that was further increased upon PDS treatment (Figure 6D). This increase occurred predominantly in S and G_2_/M phase cells. These observations suggest that HLTF prevents DNA damage when G4 structures are stabilized. Thus, although HLTF loss allows cells to tolerate G4s structures during DNA synthesis, increased DNA damage is still observed.

Finally, we evaluated the proliferation of RPE1 cells after treatment with PDS. We found that the knockdown of HLTF or MSH2 alone modestly reduced proliferation upon increasing doses of PDS, with HLTF knockdown having the greater effect. In addition, the knockdown of both HLTF and MSH2 led to a synergistic effect on proliferation (Figure 6E, F). Similar trends were observed after treatment with CX-5461, another G4 stabilizing molecule (Figure S6B, C).^92^ Collectively, these data suggest that HLTF and MSH2 act in distinct pathways to prevent DNA damage and promote cell proliferation when G4s are stabilized.

## Discussion

Here, we describe a novel function for HLTF in the maintenance of genome stability, namely in suppressing the accumulation of G4s throughout the cell cycle. HLTF can act synergistically with MSH2 in this context. *In vivo*, HLTF requires its ATPase activity to suppress G4s, but not its ubiquitin ligase or HIRAN functions, whereas all three of these activities are essential for the HLTF-mediated replication stress response. Thus, the functions of HLTF in G4 regulation and the replication stress response are distinct. Intriguingly, however, complete loss of HLTF enables tolerance to replication stress induced by G4s by suppressing fork reversal and allowing PrimPol-dependent nascent DNA synthesis. The consequence of this is increased DNA damage. Consistent with a role in preventing G4 accumulation, HLTF suppresses alternative lengthening of telomeres (ALT) activity and maintains telomere stability in an ATPase-dependent manner. Importantly, HLTF deficiency also sensitizes cells to G4 stabilization, and this is enhanced by the loss of MSH2. We propose that in cells, HLTF uses its ATP-dependent dsDNA translocase activity to suppress G4s that it encounters on DNA. Notably then, HLTF silencing, which is often observed in cancers, may have a dual effect on cells, allowing the accumulation of potentially mutagenic secondary DNA structures and allowing the replication fork to tolerate their presence.

### A function for HLTF in suppressing G4 accumulation

Several lines of evidence suggest that HLTF suppresses G4 structures in cells. First, we observe the accumulation of G4s in the absence of HLTF by imaging. Second, using ChIP-seq, we find that HLTF is enriched at G4 sites genome-wide, with its greatest enrichment at sites where multiple G4s form. Third, we observe an increase in RNA-DNA hybrids, which often co-occur with G4s, in HLTF-deficient cells. Although this increase in RNA-DNA hybrids contrasts with a previous study which reported no effect of HLTF knockdown on hybrid levels,^93^ the increase we observed is modest, in a different cell line and using a different approach for hybrid detection that may be more sensitive.^77^ Importantly, our genomic data further support the link between HLTF, RNA-DNA hybrids and G4s, as HLTF is only enriched at those hybrids that coincide with G4s. The modest increase in hybrid signal we observe in HLTF-KO cells is thus consistent with the fact that HLTF only binds a small fraction of genomic hybrids (∼15%). Notably, the concurrent formation of G4s and RNA-DNA hybrids occurs at telomeres, where each of these structures interdependently enhance the stability of the other and where their deregulation increases ALT activity.^15^ Consistent with this, we observe an increase in ALT activity in HLTF-KO cells. The localization of HLTF to telomeres in ALT-positive cells has also been reported.^94^ Altogether, our data indicate that G4s, and not RNA-DNA hybrids, drive HLTF’s localization to chromatin, and that stabilization of a G4 with a hybrid or its proximity to additional G4s further promotes this localization.

### A transcription-associated function for HLTF in the control of secondary structure accumulation

HLTF’s ability to suppress G4s appears to be distinct from its known roles in DNA damage tolerance. We found that HLTF requires its ATPase domain to suppress G4s, but not its HIRAN or ubiquitin ligase domains, which are needed for DNA damage tolerance in S phase. Indeed, G4 structures accumulate upon HLTF loss in all cell cycle phases. Also consistent with a role outside of S phase, more than half of HLTF’s chromatin binding is dependent upon transcription (Figure 1H). Since transcription unwinds duplex DNA, this process facilitates the formation of G4s and RNA-DNA hybrids. Hence, the reduction in HLTF chromatin binding following transcription inhibition may reflect the reduction of these structures and subsequent dissociation of HLTF. Alternatively, HLTF may modulate transcription and thus indirectly regulate G4 formation. Intriguingly, HLTF was originally identified as a transcription factor because it binds certain promoter motifs and positively regulates transcription in reporter assays.^42,43^ Although the mechanism underlying this function has not been well investigated, our findings support a potential role for HLTF in transcription, and it will be interesting to determine how this is linked to its regulation of secondary DNA structures.

### Proposed mechanism for HLTF-dependent regulation of G4s

Although G4s can form in ssDNA at replication forks,^88,89,95^ our data suggest that HTLF acts on G4s that form throughout the cell cycle and are associated with transcription. *In vivo*, HLTF’s suppression of G4s depends on its ATPase activity, although the precise mechanism is unclear. As ATPase activity is required for HLTF to translocate on dsDNA *in vitro*,^49,50^ one possibility is that HLTF may travel on DNA until it encounters a secondary structure which hinders its progression. HLTF’s translocase activity may then destabilize the secondary structure, allowing the structure-forming ssDNA to bind its complementary strand and restore duplex DNA. Given that G4s in transcriptionally active regions of the genome form in ssDNA, however, it is unclear if the dsDNA translocase activity will be directly required for G4 resolution. Alternatively, HLTF’s translocase activity may simply help localize it to G4s, where it may dismantle the structure or recruit another G4 resolution factor. We also speculate that resolution of RNA-DNA hybrids may be a secondary consequence of G4 resolution, consistent with the recently reported inability of HLTF to resolve R-loops *in vitro*.^93^ The enrichment of HLTF in transcriptionally active regions of the genome (Figure 2H) is consistent with our model, since G4s are often formed in transcriptionally active regions.^9^ However, it is also possible that HLTF is targeted to transcriptionally active regions of the genome to resolve G4 structures. This might allow efficient progression of RNAPII and prevent the disruption of transcription.

Another interesting question is whether HLTF can suppress the formation of other secondary DNA structures. It has been reported that HLTF can reduce the expansion of CAG/CTG or GAA/TCC repeats in human cells.^96,97^ These repeats can form DNA hairpins or H-DNA structures respectively, and their expansion is linked to the secondary structure-forming properties of the sequence. However, these studies did not identify the HLTF domain(s) required for this expansion or clearly distinguish between potential S phase or non-S phase functions of HLTF. As the analysis of many secondary structures in cells is still challenging, the question of HLTF’s role in regulating other structures will require further work.

### The synergy between HLTF and MSH2 during G4 regulation

Intriguingly, our data strongly suggest that HLTF and MSH2 have complementary roles in regulating G4s. First, we found that the chromatin binding of each protein increases when the other is lost, suggesting that each protein may compensate for the other in suppressing G4s, RNA-DNA hybrids and potentially other structures. The increase in G4s upon MSH2 knockdown was small relative to HLTF knockdown, suggesting that HLTF plays a more prominent role in regulating G4 levels than MSH2. Second, a synergistic increase in G4s was observed when both HLTF and MSH2 were absent. Third, we found that there was a further increase in cellular sensitivity to PDS when both proteins were lost. Finally, we found that both proteins act independently to suppress ALT activity. Although our findings support complementary roles for HLTF and MSH2, they do not rule out the possibility that HLTF and MSH2 may act cooperatively on some G4s. For example, MSH2 could help to melt the G4 structure, while HLTF promotes the reannealing of the melted G4 DNA with its complementary DNA strand.

Our findings also suggest HLTF and MSH2 use different mechanisms to resolve G4s. The MutSα and ꞵ complexes can directly bind to G4s and regulate their stability *in vitro*.^32,33^ *In vivo*, MutSꞵ also suppresses G4 formation.^33^ One possibility is that MutS complexes may play an indirect role in resolving G4s by the recruitment of G4 resolving helicases.^98,99^ Perhaps related, MLH1, another MMR protein acting downstream of MSH2, interacts with the G4 helicase FANCJ ^32,100^ and has been proposed to recruit it to the replication fork.^101^ Intriguingly, similar to MSH2 loss, FANCJ loss also leads to increased HLTF chromatin binding.^102^ How these observations relate to the effect we observe remains to be determined.

The complementary roles of HLTF and MSH2 in preventing the accumulation of G4s raises questions about what other factors might act with HLTF to resolve G4s. Cells have a host of factors that contribute to G4 resolution, the reasons for which are not clear. Some may be linked to the suppression of G4s that arise during different biological processes, including replication and transcription,^20,32,103^ whereas others may act on different types of G4 structures. They may also act cooperatively in the resolution process, approaching the G4 with different polarities or promoting the activity of other factors.^6,29^ The activity of certain factors may also be cell-type dependent. Indeed, we did not observe further accumulation of G4s in RPE1 cells when MSH2 was knocked down in the absence of HLTF, suggesting other factors may be more important for G4 resolution in this cell type. Regardless, when G4s are stabilized, we did observe an increased effect of their combined loss on viability indicating they have complementary roles. Clearly further work will be required to understand how and why so many factors contribute to the regulation of G4s and other secondary structures.

### HLTF and the tolerance of G4 structures

We previously found that HLTF loss promotes resistance to replication stress induced by several compounds, including HU, ATR inhibition and mitomycin C. In response to HU, we also showed that replication forks continue nascent DNA synthesis in a PrimPol-dependent manner when HLTF is depleted. This correlation led us to consider that unrestrained fork progression may underlie the observed replication stress resistance.^54^ In response to G4 ligands, however, this is not the case. Even though DNA synthesis continues in HLTF-KO cells using PrimPol, the absence of HLTF still sensitizes cells to G4 ligands, resulting in increased DNA damage and reduced proliferation. Therefore, despite the ability of cells to continue DNA replication in the presence of PDS, the stabilized G4s are still problematic for the cell and can cause DNA damage. This damage could result from the inability to resolve G4s in the post-replicative gaps and subsequent processing of these structures, and is consistent with the higher level of γH2AX observed in S and G_2_/M cells. Thus, our new data indicate that unrestrained fork progression is not sufficient for replication stress resistance in HLTF-deficient cells. As HLTF loss is associated with the transformation of cancer cells,^60^ the ability to sensitize HLTF-deficient cells to G4-stabilizing drugs could be an important observation in the context of cancers in which HLTF is silenced.

### Conclusions

In summary, our findings reveal an unexpected function for HLTF in the regulation of secondary DNA structures. Despite clear roles for HLTF in the replication stress response, HLTF helps to maintain the normal level of G4s outside of S phase and in a manner that is linked to transcription. Intriguingly, HLTF’s known functions in fork reversal and damage tolerance may synergize with its function in suppressing the formation of G4s to prevent the mutagenic and potentially toxic effects of these structures. HLTF-dependent slowing of fork progression may promote the resolution of G4s and other DNA secondary structures and prevent alternative modes of replication. Hence, HLTF loss would elevate the formation of these replication stress-inducing structures yet allow their bypass and eventual resolution by potentially mutagenic pathways. In the context of cancer, HLTF silencing could thus both elevate replication stress resulting from G4s and increase the tolerance to that stress, driving mutagenesis. In the long term, this may pose an important therapeutic opportunity for the treatment of HLTF-deficient cells with G4 stabilizing drugs.

## Acknowledgments

We thank members of the Cimprich laboratory and Gheorge Chistol for helpful discussions. We thank Miaw-Sheue Tsai for baculovirus production and insect cell expression of HLTF protein (supported by P01CA092584 from the NIH). We thank Carl Schiltz for purified proteins and for assistance with the Ub ligase assay. This work was supported by grants from the NIH to K.A.C. (ES016486 and CA092584), B.F.E. (R35GM136401), D.C. (R01GM116616 and

P01CA092584). K.A.C. is an ACS Research Professor.

## Author contributions

G.B. and K.A.C. designed the project. G.B. performed the SILAC-iPOND experiment and D.C. performed the mass spectrometry and the initial data analysis. T.E. performed the HLTF ChIP-seq experiment and G.B. performed bioinformatics analysis. M.P.C. and T.E. provided guidance on bioinformatics analysis. U.K. performed the transcription inhibition experiment. G.B. performed DNA fiber analyses with A.S. B.H.G. performed fork reversal and ATPase assays and E.M.P. performed Ub ligase assays under supervision from B.F.E. All the other experiments were performed by G.B. K.A.C. supervised the project. The manuscript was written by G.B., M.P.C., T.E., U.K. and K.A.C.

## Declaration of interests

Karlene A. Cimprich is a member of the scientific advisory board of IDEAYA Biosciences and RADD Pharmaceuticals.

## METHODS

### Cell Culture

U2OS (osteosarcoma; human female origin, ATCC #HTB-96) cells were maintained in DMEM (Life Technologies) supplemented with 10% FBS, 2 mM L-glutamine, and 100 U/mL penicillin/streptomycin. MCF10A (mammary gland epithelial; human female origin, ATCC #CRL-10317) cells were maintained in DMEM/F12 (Life Technologies) supplemented with 5% horse serum, 100 U/mL penicillin/streptomycin, 0.5μg/mL hydrocortisone, 10μg/mL insulin, 20ng/mL EGF, and 100ng/mL Cholera Toxin. p53 inactivated, Cas9 expressing hTERT-RPE1 (immortalized; retinal pigment epithelium; human female origin, ATCC #CRL-4000) cells described as RPE1 in the manuscript were received as a kind gift from Dr. Daniel Durocher and maintained in DMEM/F12 (Life Technologies) supplemented with 10% FBS, 100 U/mL penicillin/streptomycin. All cell lines were tested for the absence of mycoplasma contamination and cultured in a humidified incubator with 5% CO_2_ at 37°C.

### Construction of Cas9 expressing cells

Lentivirus encoding a construct that constitutively expresses *S. pyogenes* Cas9 was purchased (Addgene 52962-LV) and used to infect U2OS and MCF10A cells in the presence of polybrene (1 μg/mL, Millipore) overnight. 48 h post-infection, 10 μg/mL (for U2OS cells) and 5 μg/mL (for MCF10A cells) blasticidin was added to the media to select for the infected cells. Cas9 expression in the resistant cells was verified by western blotting.

### Gene expression knockdown using siRNA or sgRNA

An siRNAs smart pool for MSH2 was purchased from Dharmacon and transfected using Dharmefect 1 (Horizon) transfection reagent according to the manufacturer’s directions. siRNA against GL3 luciferease was used as a negative control. sgRNA against HLTF and MSH2 (Thermo Fisher Scientific) were transfected into Cas9 expressing U2OS, RPE1 or MCF10A cells using Lipofectamine RNAiMAX (Thermo Fisher Scientific) according to the manufacturer’s directions. A non-targeting sgRNA was used as a negative control. siRNA transfected cells were assayed 72 h post-transfection to test gene expression knockdown by western blotting, or used for other assays. sgRNA transfected RPE1 cells were assayed 72 h post-transfection, U2OS cells were assayed 96 h post-transfection to test gene expression knockdown by western blotting or immunofluorescent imaging, or used for other assays.

### Construction of rescue cell lines

HLTF’s cDNA on the pcDNA3.1(+) backbone^54^ was mutated using Q5 Site-Directed Mutagenesis kit (NEB) to generate the C760S and R890Q mutant, then cloned into pCW57.1 using HiFi assembly kit (NEB). All the plasmids were sequenced and verified. pCW57.1-HLTF vectors were packaged into lentivirus particles using the 2nd generation lentiviral packaging system (pMD2.G & psPAX2) in HEK293T cells using TransIT®-LT1 Transfection Reagent (Mirus). Virus-containing media was harvested 24 & 48h post-transfection and filtered through 0.45 μm PES membrane syringe filter to eliminate packaging cells. Lentivirus particles were further concentrated using Lenti-X Concentrator (Clontech) according to the manufacturer’s instructions. U2OS HLTF-KO cells were infected with the purified lentivirus particles in the presence of polybrene (1 μg/mL, Millipore) overnight. 48h post-infection, 1 μg/mL puromycin was added to the media to start the selection of infected cells. The resistant cells were clonally isolated using cloning cylinders. Resistant clones were further selected after doxycycline (dox) induction (500ng/mL) for 24h and HLTF expressing clones were identified by immunofluorescence staining and also verified by western blotting. Two clones for each genotype were analyzed separately and the results were later pooled and presented.

Although designed as a dox-inducible expression system, we observed that the expression of the WT, R71E and C760S mutant was similar to HLTF’s endogenous level even in the absence of dox.^54^ In experiments involving HLTF-KO cells expressing the WT and these mutants, dox was omitted. We failed to detect the expression of the R890Q mutant in HLTF-KO cells unless dox was used to induce protein expression. Therefore, in experiments evaluating HLTF-KO cells expressing the R890Q mutant, cells were either untreated (mock) or treated with dox to induce R890Q expression before assessing other phenotypes.

### iPOND-SILAC Mass Spectroscopy and Data Analysis

SILAC-labeled HEK293T cells were pulsed with 10 µM EdU with the presence of 50 µM HU for 13 min for the WT, and 10 min for the HLTF-KO cells. After labeling, roughly 6x108 asynchronous cells for each sample were fixed in 1% formaldehyde/PBS for 20 min at room temperature (RT), quenched using 0.125 M final concentration of glycine, and washed three times in PBS. Cells were scraped off the tissue culture dishes and the cell pellets were frozen at −80°C. Pellets were then resuspended in 0.25% Triton-X/PBS to permeabilize. Pellets were washed once with 0.5% BSA/PBS and once with PBS prior to the click reaction. An equal number of light and heavy SILAC-labeled cells were mixed and resuspended in a homemade click reaction buffer (10 μM biotin-PEG-azide, 2 mM CuSO_4_, 10 mM sodium L-ascorbate in PBS, freshly prepared) and incubated for 1h at RT. After cell lysis by sonication, biotinylated chromatins were captured using streptavidin-coupled C1 magnabeads for 1h at RT, then washed with lysis buffer (1% SDS in 50 mM Tris-HCl pH 8.0), low salt buffer (1% Triton X-100, 20 mM Tris-HCl pH 8.0, 2 mM EDTA, 150 mM NaCl), high salt buffer (1% Triton X-100, 20 mM Tris-HCl pH 8.0, 2 mM EDTA, 500 mM NaCl), lithium chloride wash buffer (100 mM Tris-HCl pH 8.0, 500 mM LiCl, 1% Igepal), and twice in lysis buffer. Captured proteins were eluted in SDS sample buffer by incubating for 30 min at 95°C.

iPOND samples were then processed and analyzed using mass spectrometry as described previously.^104^ For SILAC protein ratios, a minimum of two unique peptides and one or more ratio counts were required for protein group inclusion in the analysis. SILAC protein ratios for all datasets were analyzed within the Perseus software.^105^ Only proteins identified in both datasets were included in the statistical analysis in Perseus using a two-tailed t-test with a p-value set at 0.05.

### Immunofluorescent staining and imaging

Cells were seeded in 96 well optical plate and incubated overnight. The next day, 10 μM EdU diluted in cell culture media was added to label S phase cells for 30 min. After labeling, cells were pre-extracted with ice-cold CSK100 (100 mM NaCl, 300 mM sucrose, 3 mM MgCl_2_, 10 mM MOPS & 0.5% Triton X-100) buffer at 4°C for 5 min, then immediately fixed with 4% PFA/PBS for 20 min. After PBS washes, cells were permeabilized with 0.5% Triton X-100 in PBS for 5 min. A homemade click reaction buffer (10 μM biotin-PEG-azide, 2 mM CuSO_4_, 10 mM sodium L-ascorbate in PBS, freshly prepared) was added to the cells for 30 min, followed by blocking in 3% BSA/PBS for 20 min at RT. The primary antibodies were diluted in 3% BSA/PBS and incubated overnight at 4°C: rabbit anti-G4 (clone 1H6, Absolute Antibody ab00389-23.0, 1:500), goat anti-G4 (clone 1H6, Absolute Antibody ab00389-24.1, 1:500), anti-G4 (clone BG4, Sigma-Aldrich MABE917, 1:500), rabbit anti-GFP (Abcam ab290, 1:1000), rabbit anti-GFP (Thermo Fisher Scientific A-11122, 1:1000), rabbit anti-HLTF (Abcam ab183042, 1:1000), mouse anti-MSH2 (Abcam ab52266, 1:1000), mouse anti-RNA polymerase II CTD repeat YSPTSPS (phospho S2) (Abcam ab24758, 1:500), rabbit anti-RPA34 (Abcam ab97594, 1:1000). For G4 detection using BG4 antibody, rabbit anti-DYKDDDK (Cell Signalling Technology 14793S, 1:500) was diluted in 3% BSA/PBS and incubated for 1 h at RT. After primary antibody incubation, cells were washed 3x with PBS. Secondary antibodies (diluted 1:500) and DAPI (2μg/mL) were diluted in 3% BSA/PBS and incubated for 1 h at RT. Cells were washed 3x with PBS and then submerged in PBS during QIBC image acquisition.

### RNA-DNA hybrid detection by GFP-dRNH1 using fluorescent imaging

Cells were seeded in 96 well optical plate and incubated overnight. Next day, 10 μM EdU diluted in cell culture media was added to the cells to label S phase cells for 30 min. After labeling, cells were fixed with ice-cold methanol for 10 min at −20°C. RNase H diluted (1:100) in 1× RNase H buffer (New England Biolabs) was added to the cells and incubated for 2 h at 37°C. Mock-treated cells were incubated in parallel in the same buffers but without RNase H added. Following enzyme incubations, cells were washed twice in PBST (0.1% Tween 20 in PBS), then once in PBS for 5 min each. A homemade click reaction buffer (10 μM biotin-PEG-azide, 2 mM CuSO_4_, 10 mM sodium L-ascorbate in PBS, freshly prepared) was added to the cells for 30 min to detect EdU incorporated S phase cells, followed by blocking in 3% BSA/PBS for 30 min at RT. GFP-dRNH1^77^ (0.188 mg/ml stock) diluted in 3% BSA/PBS (1:2000) was added to the cells and incubated overnight at 4°C. Then, cells were washed twice in PBST and once in PBS for 5 min each. Streptavidin-conjugated Alexa 647 (diluted 1:500) and DAPI (2 μg/mL) were diluted in 3% BSA/PBS and incubated for 1 h at RT. Cells were washed 3x with PBS and then submerged in PBS during QIBC image acquisition.

### Quantitative Image-Based Cytometry (QIBC)

Images were acquired in an unbiased fashion with the Molecular Devices ImageXpress Micro automated inverted epifluorescence microscope. Acquisition times for different channels were adjusted to obtain images in non-saturating conditions for all the treatments analyzed. After acquisition, the images were analyzed with automated MetaXpress image analysis software. At least 2000 cells were analyzed per condition, and each experiment was repeated at least 3 times. DAPI signal was used for generating a mask that identified each individual nucleus as an individual object. This mask was then applied to quantify pixel intensities in the different channels for each individual cell/object. After quantification, the quantified values for each cell (mean and total intensities, area, perimeter) were extracted and exported to the proprietary Spotfire software. Spotfire was used to visualize key features of individual nuclei for thousands of cells and quantify immunofluorescent signal values at single cell level. Spotfire filtered data was then used to generate plots using GraphPad Prism [version 10.0.1 (218) for Windows 64-bit, GraphPad Software, La Jolla California USA, https://www.graphpad.com] software.

Whenever cells were labeled with EdU, total DNA content and mean EdU intensities of individual cells were used to determine cell cycle phases. Cells with positive EdU staining were considered as S phase cells. Total DNA content of the EdU negative cells were used to distinguish G_1_ (2N DNA content) from G_2_/M (4N DNA content) phase.

### ALT-associated PML body detection

Cells were seeded in 24 well optical plate and incubated overnight. The next day, cells were pre-extracted with ice-cold CSK100 (100 mM NaCl, 300 mM sucrose, 3 mM MgCl_2_, 10 mM MOPS & 0.5% Triton X-100) buffer at 4°C for 5 min, then immediately fixed with 4% PFA/PBS for 20 min. After PBS washes, cells were permeabilized with 0.5% Triton X-100 in PBS for 5 min and blocked in blocking solution (0.1% BSA, 3% goat serum, 0.1% Triton X-100, 1 mM EDTA, pH 8.0 in PBS) for 30 min at RT. The primary antibodies were diluted in 3% BSA/PBS and incubated overnight at 4°C: rabbit anti-RPA34 (Abcam ab97594, 1:1000), mouse anti-PML (Santa Cruz Biotechnology sc-966, 1:200). After primary antibody incubation, cells were washed 3x with PBS. Secondary antibodies (diluted 1:500) were diluted in 3% BSA/PBS and incubated for 1 h at RT. A second fix was performed with 2% PFA/PBS for 10 min at RT. After PBS washes, samples were then serial dehydrated with 70%, 95% and 100% ethanol for 5 min each and let air dry. Biotin-tagged TelC probe (PNA Bio F2001) was diluted (1:200) in hybridization solution [70% formamide, 1 mg/mL Blocking Reagent (Roche 11096176001), 10 mM Tris-HCl pH 7.2, in ddH_2_O]. Samples and the diluted probe were both incubated at 80°C for 5 min, before the probe was added to the sample and incubated at 80°C for another 10 min, followed by incubation at RT for 2 h. Samples were then washed twice with washing solution (70% formamide, 10 mM Tris-HCl pH 7.2, in ddH2O) and twice with PBS, 3 min each wash. Streptavidin conjugated Alexa 647 (diluted 1:500) and DAPI (2 μg/mL) were diluted in 3% BSA/PBS and incubated for 1 h at RT. Cells were washed 3x with PBS and then coverslips were mounted using ProLong Glass antifade (Thermo Fisher Scientific P36984). Random views were selected and imaged using a Zeiss OBSERVER.Z1 INVERTED microscope and a Plan-APO 40x/1.4 Oil DIC (UV) VIS-IR objective. Fluorescent images were acquired using an Axiocam 506 mono camera (conversion = 0.1135) connected to the microscope. Images were analyzed using ImageJ2 Fiji to count the number of PML and telomere FISH colocalizaing foci per cell.

### In situ single-stranded telomeric C-strand (ssTelo-C) detection

Cells were seeded in a 24 well optical plate and incubated overnight. The next day, cells were fixed with 4% PFA/PBS for 20 min. ssTelo-C was stained as described previously.^87^ Briefly, samples were incubated with 500 μg/mL RNase A (Thermo Fisher Scientific EN0531) diluted in blocking solution (0.1% BSA, 3% goat serum, 0.1% Triton X-100, 1 mM EDTA, pH 8.0 in PBS) for 1h at 37°C. Samples were then washed once with PBS, then serial dehydrated with 70%, 95% and 100% ethanol for 5 min each and let air dry. FAM-tagged TelG probe (PNA Bio F1005) was diluted (1:100) in hybridization solution [70% formamide, 1 mg/mL Blocking Reagent (Roche 11096176001), 10 mM Tris-HCl pH 7.2, in ddH_2_O] and added to the samples to incubate for 2 h at RT. Samples were washed twice with washing solution (70% formamide, 10 mM Tris-HCl pH 7.2, in ddH2O) and twice with PBS, 3 min each wash. Nuclei were counterstained with DAPI (2 μg/mL) diluted in PBS and incubated for 1 h at RT. Cells were washed 3x with PBS and then coverslips were mounted using ProLong Glass antifade (Thermo Fisher Scientific P36984). Random views were selected and imaged using a Zeiss OBSERVER.Z1 INVERTED microscope and a Plan-APO 40x/1.4 Oil DIC (UV) VIS-IR objective. Fluorescent images were acquired using an Axiocam 506 mono camera (conversion = 0.1135) connected to the microscope. Images were analyzed using ImageJ2 Fiji to count the number of single-stranded TelC foci per cell.

### ChIP-seq with spike-in normalization (ChIP-Rx)

For each ChIP-Rx sequencing experiment, 6x10^7^ cells per immunoprecipitation condition were fixed with 1% formaldehyde (Fisher Chemical, F79-500) for 5 min at RT. Fixation was stopped by adding 125 mM glycine for 5 min. Cells were harvested in ice-cold PBS containing protease (Sigma-Aldrich, 11836170001) and phosphatase inhibitors (Roche, 4906837001). All further used buffers also contained these inhibitors. As exogenous control (spike-in), murine ESC cells expressing MPP8-GFP (gift from Wysocka Lab ^106^) were added at a 1:10 ratio during cell lysis. Cell lysis was carried out for 20 min in lysis buffer I (5 mM PIPES pH 8.0, 85 mM KCl, 0.5% NP-40 in nuclease-free water) and nuclei were collected by centrifugation at 500 g for 20 min at 4°C. Crosslinked chromatin was prepared in lysis buffer II (10 mM Tris-HCl pH 7.5, 150 mM NaCl, 1 mM EDTA, 1% NP-40, 1% sodium deoxycholate, 0.1% SDS in nuclease-free water) and fragmented by using the Covaris Focused Ultrasonicator E220 for 20 min (10% duty factor, 140 PIP, 200 CPB) per mL lysate. Fragment size of 150-300 bp was validated by agarose gel electrophoresis. Chromatin was centrifuged for 20 min at 14,000 rpm at 4 °C before IP. For each IP reaction, 100 μl Dynabeads Protein A and Protein G (Thermo Fisher Scientific, 10002D and 10004D) were pre-incubated overnight with rotation in the presence of 5 mg/ml BSA/PBS (Thermo Fisher Scientific, BP1600-100) and 15 μg antibody GFP (Abcam, ab290). Chromatin was added to the beads, and IP was performed for at least 6 h at 4 °C with rotation. Beads were washed three times each with washing buffer I (20 mM Tris-HCl pH 8.1, 150 mM NaCl, 2 mM EDTA, 1% Triton X-100, 0.1% SDS in nuclease-free water), washing buffer II (20 mM Tris-HCl pH 8.1, 500 mM NaCl, 2 mM EDTA, 1% Triton X-100, 0.1% SDS in nuclease-free water), washing buffer III (10 mM Tris-HCl pH 8.1, 250 mM LiCl, 1 mM EDTA, 1% NP-40, 1% sodium deoxycholate in nuclease-free water), and once with TE buffer (Invitrogen). Chromatin was eluted twice by incubating with 150 mL elution buffer (100 mM NaHCO3, 1% SDS) for 15 min with rotation at RT. Input samples and eluted samples were de-crosslinked overnight at 65 °C. Protein and RNA were digested with proteinase K (Thermo Scientific, 25530049) and RNase A (Thermo Scientific, EN0531), respectively. DNA was isolated by phenol-chloroform extraction and ethanol precipitation and analyzed by qPCR using Light Cycler Real-Time PCR System (Roche) and iTaq Universal SYBR Green Supermix SYBR Green Master Mix (BioRad, 1725122) before library preparation followed by high throughput sequencing on a Novaseq 6000.

For ChIP-Rx sequencing, DNA was quantified using the Quant-iT™ dsDNA HS Assay Kit (Invitrogen, Q32851). DNA library was prepared using the NEBnext Ultra II DNA Library Prep Kit (New England Biolabs, E7645) and NEBNext Multiplex Oligos for Illumina (NEB, E7600) following the manufacturer’s instructions. Cycle number was determined by qPCR (Thermo Scientific, S11494). Quality and Quantity of the library was assessed on the QUBIT using the Quant-iT™ dsDNA HS Assay Kit, the 2100 Bioanalyzer (Agilent) using the High Sensitivity DNA Assay and NEBNext Library Quant Kit for Illumina (NEB, E7630). Finally, libraries were subjected to cluster generation and base calling for 150 cycles paired-end on the Illumina Novaseq 6000 platform.

### Bioinformatics analysis

For HLTF ChIP-seq samples, FASTA files of human (hg38) and mouse (mm39) reference genome were downloaded from RefSeq (https://www.ncbi.nlm.nih.gov/datasets/genome/). Both FASTA files were used in bowtie2-build to generate an indexed “hybrid” reference genome. Raw FASTQ reads from HLTF ChIP-seq were aligned to the hybrid reference genome using bowtie2^107^ (2.2.5) –local mode using paired inputs with default parameters. Duplicates were identified using Picard (2.27.5) MarkDuplicates. Samtools (1.16.1) was used to filtered the reads: reads aligned to the chromosomes were kept; unmapped, pairs with only one mate aligned, duplicates and reads aligned to the ENCODE blacklist regions were excluded.^108^ After filtering, the reads are parsed into human versus mouse aligned reads. The total number of mouse aligned reads for each sample were taken to calculate the spike-in normalization factors as previously described.^72^ Deeptools^109^ (3.5.1) bamCoverage was used to generate bigwig files with the signals scaled using the spike-in normalization factor. The resulting bigwig files were used in deeptools computeMatrix, followed by plotHeatmap and plotProfiles to generate the heatmaps and aggregate plots respectively. For G4 ChIP-seq samples (GSE162299), raw reads in FASTQ files were downloaded and aligned to human (hg38) reference genome using bowtie2 (2.2.5) –local mode with default parameters. The aligned reads were deduplicated and filtered as mentioned above. Deeptools (3.5.1) bamCoverage was used to generate bigwig files with the signals normalized to 1x human genome size, using the reads per genome content (RPGC) method. Bed files of the G4 CUT&Tag peaks (GSE181373), DRIP peaks (GSE115957) identified in the U2OS cells and *in vitro* G4 forming motifs in the human genome under PDS^+^, K^+^ condition (GSE110582) were downloaded and used to calculate HLTF ChIP-seq signal enrichment. Liftover (https://genome.ucsc.edu/cgi-bin/hgLiftOver) was used to convert genome coordinates to those in the hg38 assemblies. MACS2^110^ (2.2.7.1) was used to identify HLTF ChIP-seq peaks with default settings using FDR=0.05 as cutoff. The peaks were further filtered based on the average ChIP-seq signal intensity within the peak regions following ENCODE guidelines ^111^: at least a 2-fold signal enrichment in the control versus HLTF knockdown ChIP-seq sample was used at cut-off. Bedtools^112^ (2.30.0) was used to intersect G4 CUT&Tag, G4 motifs, DRIP and HLTF ChIP-seq peaks. HOMER^113^ (4.11) was used to analyze the basic genome ontology of G4 CUT&Tag and HLTF ChIP-seq peaks.

### DNA fiber analysis with triple nucleotide labeling

Cells were seeded in 24 well plate and incubated overnight. Unless otherwise stated in the figure legends, on the next day, cells were labeled consecutively with 50 μM IdU, 100 μM EdU and 200 μM CldU for 30, 30 and 20 min each. Warm PBS was applied to wash cells between each labeling. Drug or mock treatment was performed during the EdU labeling. After the CldU labeling, cells were trypsinized and harvested. Chromatin fibers were spread on glass slides and then fixed with freshly prepared fixative (75% methanol, 25% acetic acid) for 10 min at RT. To detect nucleotide analogue labeled fibers, slides were denatured with 2.5N HCl for 30 min at RT. A homemade click reaction buffer (100 μM biotin-PEG-azide, 2 mM CuSO_4_, 10 mM sodium L-ascorbate in PBST, freshly prepared) was added to the slides and incubated for 30 min at RT. Primary antibodies were diluted in 3% BSA/PBST (0.1% Triton X-100 in PBS) and incubated with the samples at 37°C for 1 h: mouse anti-BrdU (BD Biosciences 347580, 1:75, for IdU detection), rat anti-BrdU (Abcam ab6326, 1:100, for CldU detection), rabbit anti-biotin (Cell Signaling Technology 5597, 1:100, for biotinylated EdU detection after click reaction). PBST was used to wash the slides 3 times. Secondary antibodies diluted in 3% BSA/PBST were incubated with the samples at 37°C for 1 h: goat anti-rat IgG (H+L) (Thermo Fisher Scientific A11006, 1:100), goat anti-rabbit IgG (H+L) (Thermo Fisher Scientific A21244, 1:100), goat anti-mouse IgG1 (Thermo Fisher Scientific A21124, 1:100). Samples were then washed with PBST and let air dry, before mounted with Fluoroshield (Sigma-Aldrich F6182) using 1.5N coverglass. Random views were selected and imaged using a Zeiss OBSERVER.Z1 INVERTED microscope and a Plan-APO 40x/1.4 Oil DIC (UV) VIS-IR objective. Fluorescent images were acquired using an Axiocam 506 mono camera (conversion = 0.1135) connected to the microscope. To evaluate the replication fork progression rate, EdU tract lengths were measured only when preceded by IdU labeling and followed by CldU labeling. For quantification, at least 2 slides per sample were prepared for each experimental repeat. To avoid bias, after immunodetection, each pair of slides was blinded and we randomly selected at least 5 fields of view from each slide and acquired images. Hence, for each experiment, we acquired at least 10 images for each sample. DNA fiber length was measured using an ImageJ plug-in. We randomly scored a similar number of fibers (∼15) from each image. Measurements from at least 3 repeats were pooled, resulting in at least 200 replication tracts per sample.

### Protein purification

Human HLTF proteins^53^ and UBC13/MMS2 were expressed and purified as described previously.^114^ UBA1 (ab271808) and ubiquitin (ab269109) were purchased from Abcam.

### ATPase Assay

The ATP hydrolysis rate of HLTF was determined using a NADH-coupled enzymatic assay.^115^ Briefly, the reactions were carried out in solution containing 20 mM Tris-HCl pH 7.76, 2 mM MgCl_2_, 10 mM KCl, 1 mM DTT, 2 mM ATP, 3 mM phosphoenolpyruvate, 0.044 mM NADH, 12 U pyruvate kinase, and 17 U lactate dehydrogenase, supplemented with 100 nM HLTF, and 300 nM fork DNA (assembled from oligos 40, F20.40, LEAD20.20, and LAG20.20; see STAR method). The A340 was monitored for 60 min (1 min intervals) at 37°C in a 96-well plate (Corning) using a Synergy H1 Hybrid Reader (Biotek). Data were analyzed using GraphPad Prism and rates were corrected for background NADH decomposition from a no-enzyme control.

### Fork Regression

Fork regression was performed as previously described.^53^ Reactions containing 5 nM HLTF and 1 nM ^32^P-labeled fork DNA assembled from oligos 48, 50, 52, and 53 (see STAR method) were incubated at 37°C in reaction buffer (20 mM Tris-HCl pH 7.76, 2 mM MgCl_2_, 10 mM KCl, 1 mM DTT, 2 mM ATP, 100 µg/ml BSA).

### Ubiquitin Ligase

DNA oligonucleotides Bio-31 and Bio-75 (STAR method) were annealed in annealing buffer (10 mM Tris pH 7.5, 50 mM NaCl, 1 mM EDTA) to create a duplex with a 5’-ssDNA overhang. Reactions were carried out in ubiquitylation buffer (40 mM Tris pH 7.76, 8 mM MgCl_2_, 10% glycerol, 50 mM NaCl) and contained 0.1 μM UBA1, 40 μM ubiquitin, 0.2 μM UBC13/MMS2, 0.2 μM HLTF, 0.05 μM Bio-31/Bio-75 DNA, and 0.5 mM ATP. Reactions were incubated at 30°C for 15 minutes and stopped by the addition of 2X Laemmli buffer followed by incubation at 70°C for 5 minutes. Samples were analyzed by western blot using anti-ubiquitin antibody (P4D1, Cytoskeleton AUB01-HRP, 1:1000) and visualized with Clarity Western ECL Substrate (BioRad).

### Drug dose response assay

200 RPE1 cells were seeded per well in a 96 well plate. After growth for 24 h, PDS was added at the indicated concentration. To monitor proliferation, cells were grown in an IncuCyte Zoom live cell imager (Essen/Sartorius) with images recorded every 4 h for 6 d after addition of the drug. PDS containing media was refreshed 3 d after initial drug addition. The confluence percentage for each image was calculated using the IncuCyte Zoom software over the course of the experiment for each well. For viability measurement, the confluence level at each time point was first normalized to the initial confluence level of the cells before the addition of PDS to account for different initial cell plating density. About 5 d post drug addition, the relative cell proliferation is calculated by dividing the normalized growth of drug treated cells to that of untreated cells. The resulting relative percentage of proliferation and the corresponding drug concentration were fitted to a 4-parameter dose-response curve to calculate IC50 values.

### Western blot analysis

Cells were collected and resuspended in lysis buffer [50 mM Tris-HCl pH 7.5, 150 mM NaCl, 1 mM EDTA, 1% Triton X-100, supplemented with 1x protease inhibitor cocktail (Sigma 11836170001)]. After vortexing at 4°C for 15 min, the lysate was further sonicated using a probe sonicator. Supernatant was collected after high speed centrifugation. Protein concentration was determined using BCA assay (Thermo Scientific 23227). Equal amounts of total proteins were boiled in Laemmli sample buffer supplemented with 5% beta-mercaptoethanol for 5 min. Proteins were separated by SDS-PAGE, and transferred to a PVDF membrane (Millipore IEVH85R). Primary antibodies were: rabbit anti-HLTF (Abcam ab183042 1:5000), mouse anti-MSH2 (abcam ab52266 1:1000), mouse anti-alpha-tubulin (Sigma T9026 1:5000). Secondary antibodies were goat anti-rabbit HRP (Molecular Probes G21234) and goat anti-mouse HRP (Invitrogen 81-6520). Chemi-luminescence was carried out using the Immobilon HRP substrate (Millipore WBKLS0500), and blots were imaged with a Azure cSeries (300) Imager from Azure Biosystems.

## QUANTIFICATION AND STATISTICAL ANALYSIS

All experiments, if not indicated otherwise in the figure legend, were performed three times and representative experiments were depicted. No statistical methods or criteria were used to estimate sample size or to include/exclude samples.

Statistical analysis was performed using Prism (GraphPad Software). Details of how data is presented, including the definition of center (mean or median) and error bars can be found in the figure legends. Details of statistical test for each experiment, including the type of statistical tests used and the number of repeats, can be found in the figure legends. Statistical test results, presented as levels of significance, are shown in the figures. In all cases: ns, not significant; ∗ p < 0.05; ∗∗ p < 0.01; ∗∗∗ p < 0.001; ∗∗∗∗ p < 0.0001

## Supplemental figures

**Figure S1.**
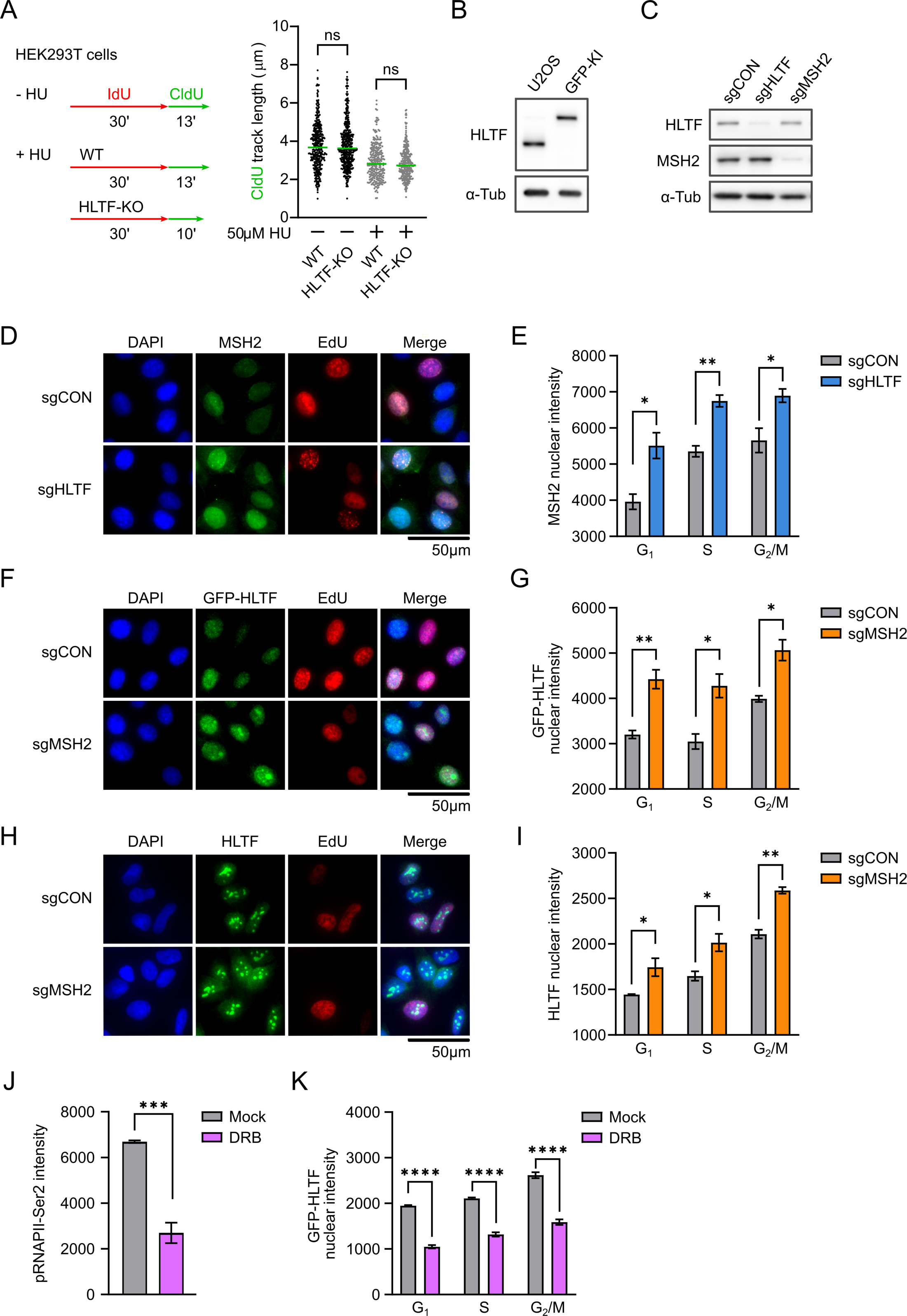
The cell cycle independent regulation of HLTF chromatin binding by MSH2 and transcription. Related to Figure 1. A. Left: experiment setup of replication fork progression assay. Right: dot plot of CldU tract lengths (n = 3). Line represents median. ns, not significant, by Mann-Whitney test. B. Western blot analysis of WT and GFP-HLTF U2OS cells. An HLTF antibody was used for HLTF detection. C. Western blot in GFP-HLTF U2OS cells 96h after sgRNA transfection. D. Representative IF images in U2OS cells expressing GFP-HLTF 96h after sgRNA transfection. E. Bar graph showing the chromatin-bound mean MSH2 intensity in U2OS cells expressing GFP-HLTF, as shown in D. The median of the mean G4 intensity is used to calculate the mean ± SEM (n=3). ∗ p < 0.05; ∗∗ p < 0.01, by t-test. F. Representative IF images in U2OS cells expressing GFP-HLTF 96h after sgRNA transfection. A GFP antibody was used to detect GFP-HLTF. G. Bar graph showing the chromatin-bound mean GFP-HLTF intensity in U2OS cells expressing GFP-HLTF, as shown in F. The median of the mean G4 intensity is used to calculate the mean ± SEM (n=3). ∗ p < 0.05; ∗∗ p < 0.01, by t-test. H. Representative IF images in U2OS cells 96h after sgRNA transfection. An HLTF antibody was used to detect HLTF. I. Bar graph showing the chromatin-bound mean HLTF intensity in U2OS cells, as shown in H. The median of the mean G4 intensity is used to calculate the mean ± SEM (n=3). ∗ p < 0.05; ∗∗ p < 0.01, by t-test. J. Bar graph showing the phosphorylated Ser2 of RNA polymerase II C-terminal domain levels (pRNAPII-Ser2) in U2OS cells expressing GFP-HLTF after 4h of mock or DRB (100 µM) treatment. The median of the mean pRNAPII-Ser2 intensity is used to calculate the mean ± SEM (n=4). ∗∗∗ p < 0.001, by t-test. K. Bar graph showing the chromatin-bound mean HLTF intensity in each cell cycle phase in U2OS cells expressing GFP-HLTF after 4h of mock or DRB (100 µM) treatment. A GFP antibody was used for IF detection of GFP-HLTF. The median of the mean GFP-HLTF intensity is used to calculate the mean ± SEM (n=4). ∗∗∗∗ p < 0.0001, by t-test.

**Figure S2.**
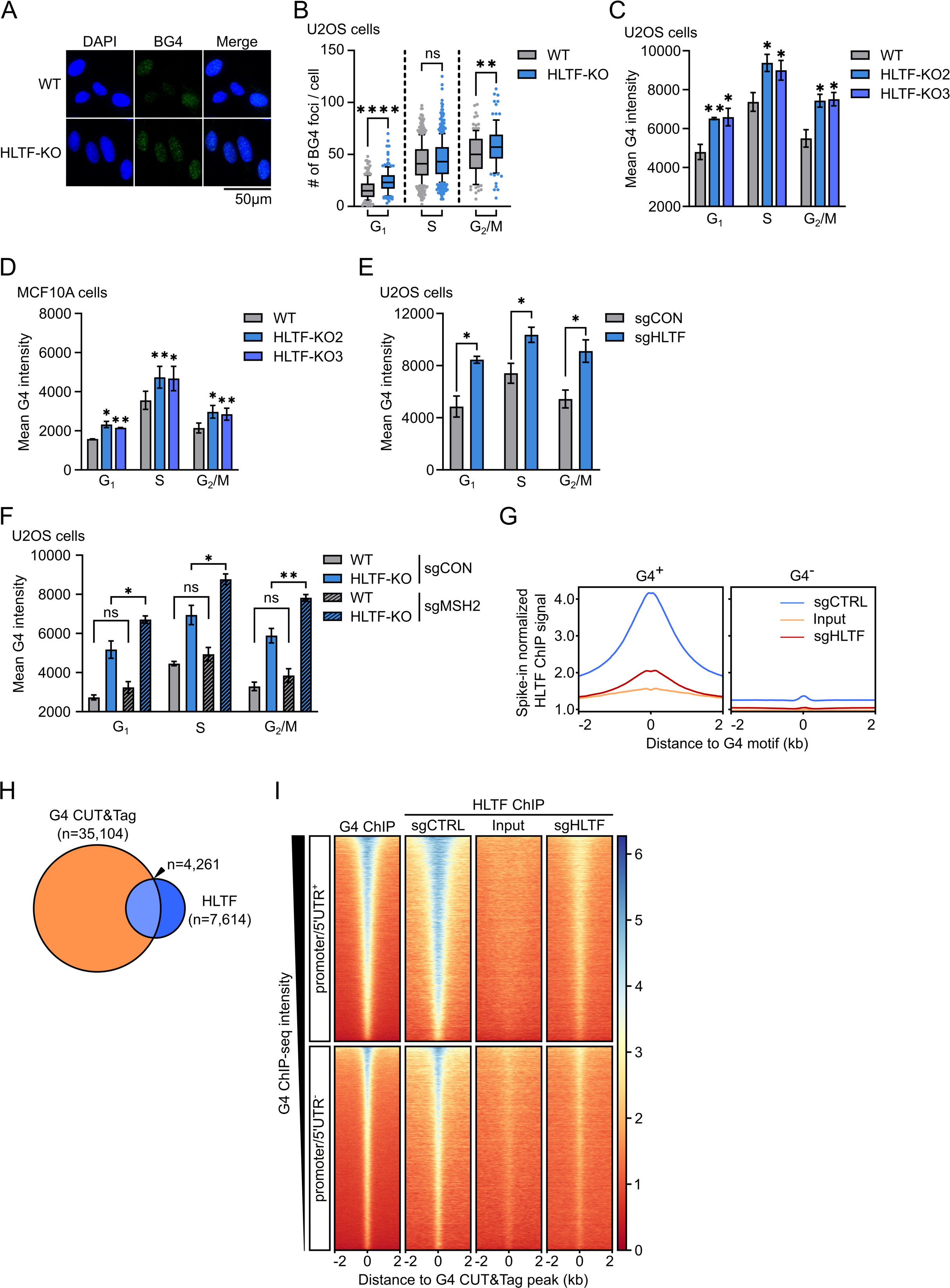
HLTF suppresses G4 accumulation in the cells and is enriched at G4 structures in the human genome. **Related to** Figure 2. A. Representative IF images in WT and HLTF-KO isogenic U2OS cells. G4 is detected using the BG4 antibody. B. Box plot showing the average number of G4 foci in WT and HLTF-KO isogenic U2OS cells, as shown in A. Whiskers indicate the 10th and 90th percentiles, boxes span the 25th to 75th percentiles. Lines inside boxes represent medians. ∗∗ p<0.01; ∗∗∗∗ p<0.0001; ns, not significant, by Mann-Whitney test. C. Bar graph showing the mean G4 intensity in WT and HLTF-KO isogenic U2OS cells. The median of the mean G4 intensity is used to calculate the mean ± SEM (n=3). Statistical test results compared to the WT cells are shown. ∗ p < 0.05; ∗∗ p < 0.01, by one-way ANOVA followed by Dunnett’s test. D. Bar graph showing the mean G4 intensity in WT and HLTF-KO isogenic MCF10A cells. The median of the mean G4 intensity is used to calculate the mean ± SEM (n=3). Statistical test results compared to the WT cells are shown. ∗ p < 0.05; ∗∗ p < 0.01; ∗∗∗ p < 0.001, by one-way ANOVA and then Dunnett’s test. E. Bar graph showing the mean G4 intensity in U2OS cells after sgRNA transfection. The median of the mean G4 intensity is used to calculate the mean ± SEM (n=3). ∗ p < 0.05, by t-test. F. Bar graph showing the mean G4 intensity in WT and HLTF-KO isogenic U2OS cells after sgRNA transfection. The median of the mean G4 intensity is used to calculate the mean ± SEM (n=3). ∗ p < 0.05; ∗∗ p < 0.01; ns, not significant, by t-test. G. Aggregate plot showing HLTF ChIP-seq coverage (y-axis) relative to the distance (x-axis) from G4 motifs (GSE110582) identified in the human genome *in vitro* in kb. G4 motifs are segregated based on whether they are within G4 structures identified in U2OS cells by G4 CUT&Tag: G4 motifs found in G4 CUT&Tag peaks are G4^+^ (n=55,768) or otherwise G4^-^ (n=1,207,711). H. Venn diagram showing the number of G4 CUT&Tag, HLTF ChIP-seq, and overlapping peaks. I. Heatmaps showing ChIP-seq coverage for G4 and HLTF at G4 CUT&Tag peaks. The x-axis represents the distance from the G4 CUT&Tag peak in kb. G4 peaks are segregated based on whether they are within the promoter or 5’UTR. Promoters are identified as ±1kb of the annotated TSS. Heatmaps were sorted by G4 ChIP-seq signal intensity for all ChIP-seq samples. Spearman correlation coefficient between G4 and HLTF ChIP-seq signal (sgCON) was 0.64 for G4 peaks in promoter or 5’UTR (promoter/5’UTR^+^), and 0.70 for G4 peaks out of promoter or 5’UTR (promoter/5’UTR^-^). Related to Figure 2I.

**Figure S3.**
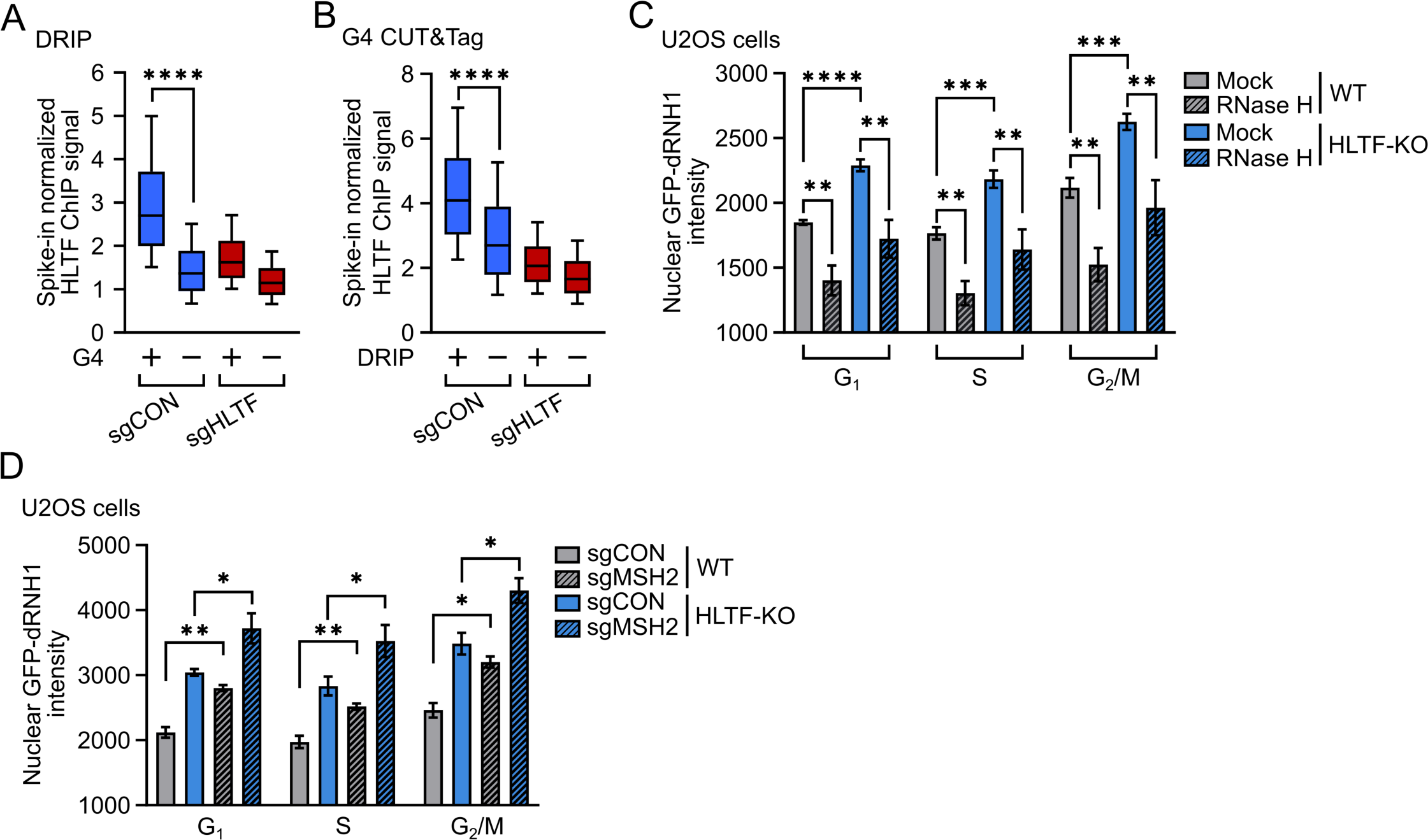
HLTF is enriched at G4s stabilized by RNA-DNA hybrids. Related to Figure 3. A. Box plot showing mean HLTF ChIP-seq signal intensity in all RNA-DNA hybrid peaks identified by DRIP-seq in U2OS cells. DRIP peaks are segregated based on whether they overlap with G4 peaks identified by G4 CUT&Tag: DRIP peaks that overlap with G4 peaks are G4^+^ (n=11,499) or otherwise G4^-^ (n=59,090). ∗∗∗∗ p<0.0001, by t-test. B. Box plot showing mean HLTF ChIP-seq signal intensity in all G4 CUT&Tag peaks identified in U2OS cells. G4 peaks are segregated based on if they overlap with RNA-DNA hybrid peaks identified by DRIP-seq: G4 peaks that overlap with DRIP peaks are DRIP^+^ (n=10,792) or otherwise DRIP^-^ (n=24,312). ∗∗∗∗ p<0.0001, by t-test. Whiskers indicate the 10th and 90th percentiles, boxes span the 25th to 75th percentiles. Lines inside boxes represent medians. C. Bar graph showing the mean nuclear RNA-DNA hybrid intensity in WT and HLTF-KO isogenic U2OS cells after RNase H digestion. The median of the mean nuclear RNA-DNA hybrid intensity is used to calculate the mean ± SEM (n=3). ∗∗ p< 0.01; ∗∗∗ p< 0.001; ∗∗∗∗ p< 0.0001, by t-test. D. Bar graph showing the mean nuclear RNA-DNA hybrid intensity in each cell cycle phase in WT and HLTF-KO isogenic U2OS cells after sgRNA transfection. The median of the mean nuclear RNA-DNA hybrid intensity is used to calculate the mean ± SEM (n=3). ∗ p< 0.05; ∗∗ p< 0.01, by t-test.

**Figure S4.**
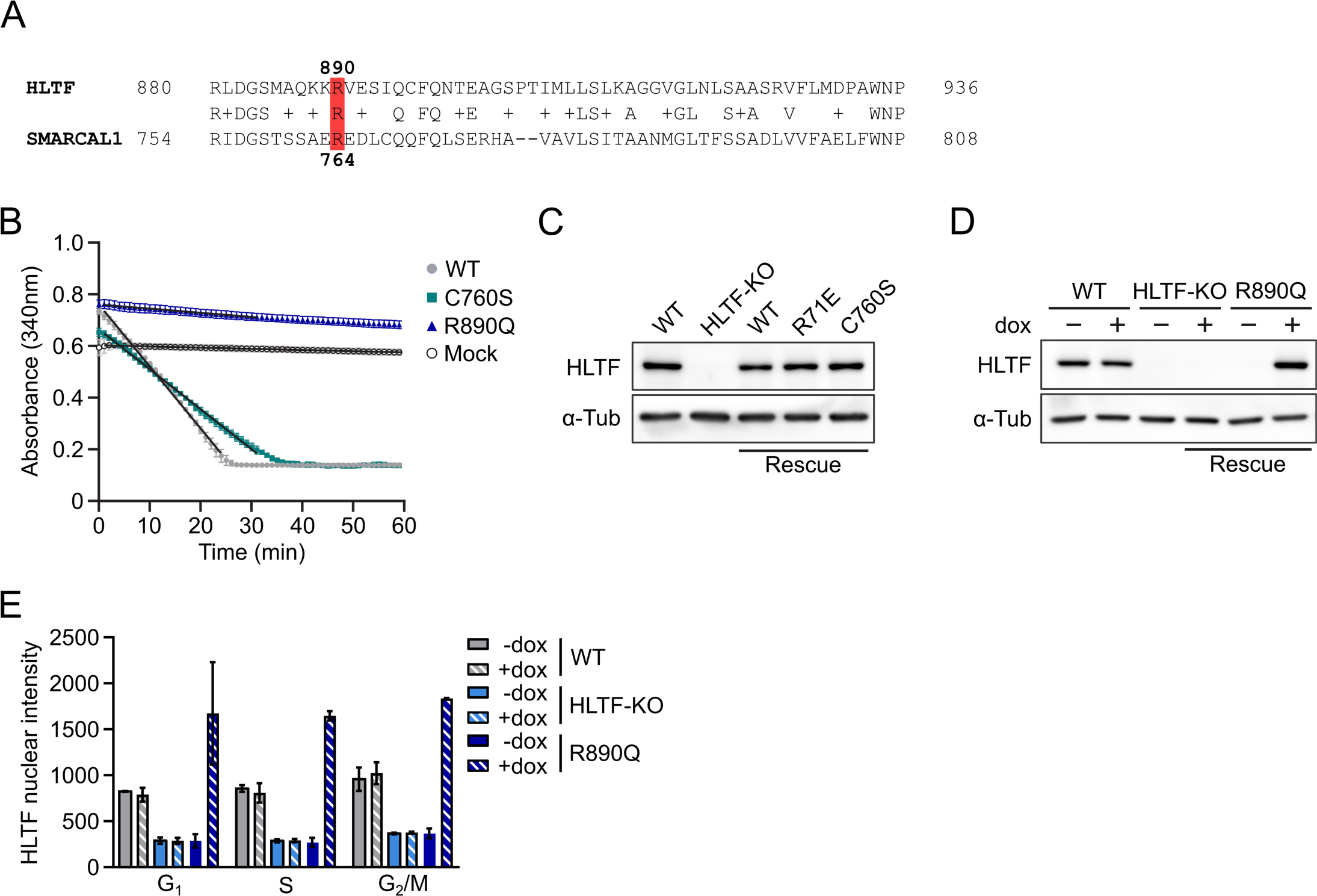
HLTF suppresses G4s in an ATPase-dependent manner in cells. Related to Figure 4. A. Protein sequence alignment of HLTF and SMARCAL1 at their C-terminus. HLTF’s R890 and SMARCAL1’s R764 residues are highlighted in red. B. Dot and line graph showing the measurement of NADH-coupled regenerative ATPase activity. Slopes used to calculate the ATPase rates shown in Figure 4B are shown as black lines and were determined by linear least squares regression of the linear range of each trace. Mock, no-enzyme control used to calculate background NADH decomposition. Data are plotted as the mean ± SD (n=3). C. Western blot in WT and HLTF-KO isogenic U2OS cells expressing HLTF WT or mutant proteins. D. Western blot in WT and HLTF-KO isogenic U2OS cells expressing the HLTF R890Q mutant, after 24 h of dox (500 ng/mL) induction. Note that no R890Q mutant expression was observed in the absence of dox. See method details. E. Bar graph showing chromatin-bound HLTF levels in WT and HLTF-KO isogenic U2OS cells, and HLTF-KO cells expressing the R890Q mutant, after 24 h of dox (500 ng/mL) induction.

**Figure S5.**
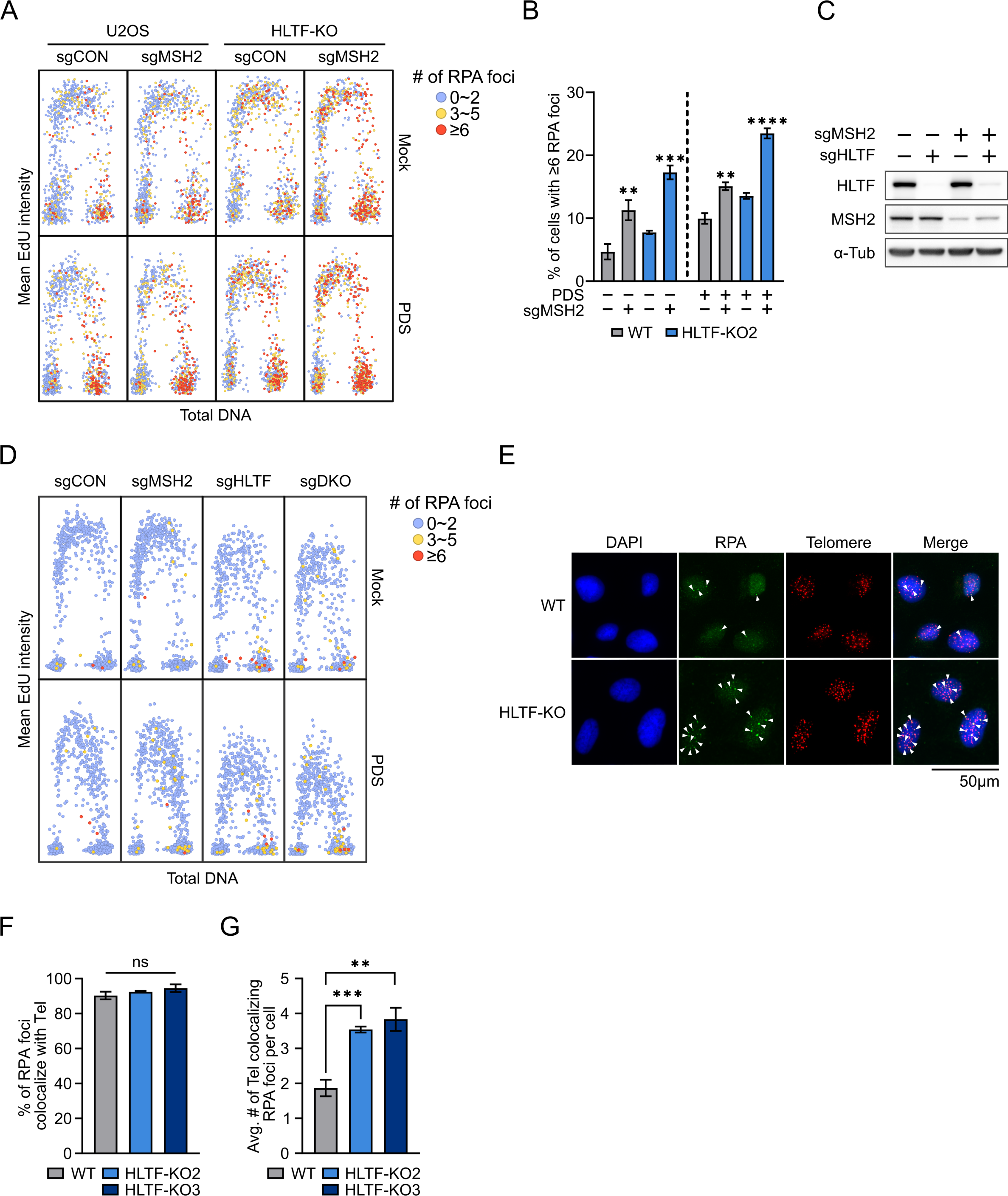
HLTF suppresses ALT activity in an ATPase-dependent manner. Related to Figure 5. A. Scatter plots of total DNA content versus mean EdU intensity in U2OS cells after sgRNA transfection and 24 h of PDS (10 μM) treatment. The number of large RPA foci per cell is shown by a color scale. 1000 cells are shown for each condition. Results from one representative experiment are shown. B. Bar graph showing mean percentage of cells with at least 6 large RPA foci, related to A. Data are represented as mean ± SEM (n = 4). One-way ANOVA followed by Dunnett’s test was performed comparing 4 mock treated samples, or 4 PDS treated samples. Test results comparing to the WT U2OS mock or PDS treated samples were shown. ∗∗ p < 0.01; ∗∗∗ p < 0.001; ∗∗∗∗ p < 0.0001. C. Western blot in RPE1 cells 72 h after sgRNA transfection. D. Scatter plots of total DNA content versus mean EdU intensity in RPE1 cells after sgRNA transfection and 24 h of PDS (5 μM) treatment. The number of large RPA foci per cell is shown by a color scale. 1000 cells are shown for each condition. Results from one representative experiment are shown. E. Representative IF images of large RPA foci and telomere detection in WT and HLTF-KO isogenic U2OS cells. Arrowheads mark the telomere colocalizing large RPA foci. F. Bar graph showing the percentage of large RPA foci that colocalize with telomeres in WT and HLTF-KO isogenic U2OS cells. Data are represented as mean ± SEM (n=3). ns, not significant by one-way ANOVA. G. Bar graph showing the average number of large RPA foci colocalizing with telomeres per cell in WT and HLTF-KO isogenic U2OS cells. Data are represented as mean ± SEM (n=4). ∗∗ p < 0.01; ∗∗∗ p < 0.001 by one-way ANOVA followed by Dunnett’s test.

**Figure S6.**
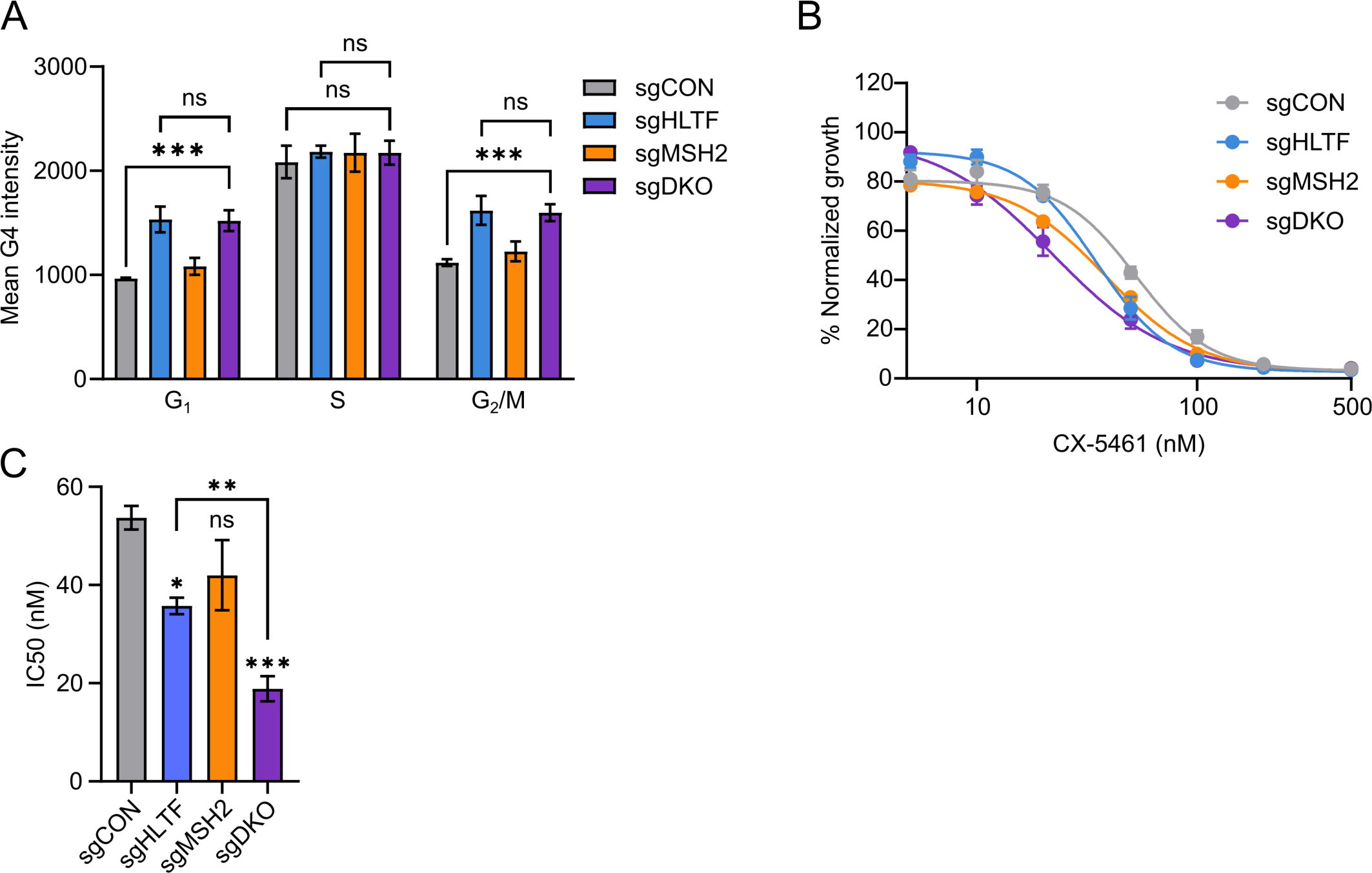
HLTF restrains replication fork progression, protects cells from DNA damage and growth defects in response to G4-stabilization. **Related to** Figure 6. A. Bar graph showing the mean G4 intensity in each cell cycle phase in RPE1 cells, 72 h after sgRNA transfection. The median of the mean G4 intensity is used to calculate the mean ± SEM (n=5). Cells are transfected with sgRNAs against both HLTF and MSH2 in the sgDKO sample. ∗∗∗ p < 0.001; ns, not significant, by t-test. B. Drug-response curve of CX-5461 in RPE1 cells after sgRNA transfection, using confluence as a readout after six days of treatment. Cells are transfected with sgRNAs against both HLTF and MSH2 in the sgDKO sample. Data are represented as mean ± SEM (n=4). C. Bar graph showing the IC50 value of RPE1 cells in response to CX-5461 treatment after sgRNA transfection, as shown in B. Data are represented as mean ± SEM (n=4). ∗ p < 0.05; ∗∗∗ p < 0.001; ns, not significant by one-way ANOVA followed by Dunnett’s test compared to sgCON. ∗∗ p < 0.01, by t-test.

## Reference

1. Mirkin, E.V., and Mirkin, S.M. (2007). Replication fork stalling at natural impediments. Microbiol. Mol. Biol. Rev. 71, 13–35.

2. Khristich, A.N., and Mirkin, S.M. (2020). On the wrong DNA track: Molecular mechanisms of repeat-mediated genome instability. J. Biol. Chem. 295, 4134–4170.

3. Brown, R.E., and Freudenreich, C.H. (2021). Structure-forming repeats and their impact on genome stability. Curr. Opin. Genet. Dev. 67, 41–51.

4. Wang, G., and Vasquez, K.M. (2023). Dynamic alternative DNA structures in biology and disease. Nat. Rev. Genet. 24, 211–234.

5. Varshney, D., Spiegel, J., Zyner, K., Tannahill, D., and Balasubramanian, S. (2020). The regulation and functions of DNA and RNA G-quadruplexes. Nat. Rev. Mol. Cell Biol. 21, 459–474.

6. Sato, K., and Knipscheer, P. (2023). G-quadruplex resolution: From molecular mechanisms to physiological relevance. DNA Repair 130, 103552.

7. Gellert, M., Lipsett, M.N., and Davies, D.R. (1962). Helix formation by guanylic acid. Proc. Natl. Acad. Sci. U. S. A. 48, 2013–2018.

8. Sen, D., and Gilbert, W. (1988). Formation of parallel four-stranded complexes by guanine-rich motifs in DNA and its implications for meiosis. Nature 334, 364–366.

9. Hänsel-Hertsch, R., Beraldi, D., Lensing, S.V., Marsico, G., Zyner, K., Parry, A., Di Antonio, M., Pike, J., Kimura, H., Narita, M., et al. (2016). G-quadruplex structures mark human regulatory chromatin. Nat. Genet. 48, 1267–1272.

10. Spiegel, J., Cuesta, S.M., Adhikari, S., Hänsel-Hertsch, R., Tannahill, D., and Balasubramanian, S. (2021). G-quadruplexes are transcription factor binding hubs in human chromatin. Genome Biol. 22, 117.

11. Shen, J., Varshney, D., Simeone, A., Zhang, X., Adhikari, S., Tannahill, D., and Balasubramanian, S. (2021). Promoter G-quadruplex folding precedes transcription and is controlled by chromatin. Genome Biol. 22, 143.

12. Vannier, J.-B., Pavicic-Kaltenbrunner, V., Petalcorin, M.I.R., Ding, H., and Boulton, S.J. (2012). RTEL1 dismantles T loops and counteracts telomeric G4-DNA to maintain telomere integrity. Cell 149, 795–806.

13. Moye, A.L., Porter, K.C., Cohen, S.B., Phan, T., Zyner, K.G., Sasaki, N., Lovrecz, G.O., Beck, J.L., and Bryan, T.M. (2015). Telomeric G-quadruplexes are a substrate and site of localization for human telomerase. Nat. Commun. 6, 7643.

14. Jansson, L.I., Hentschel, J., Parks, J.W., Chang, T.R., Lu, C., Baral, R., Bagshaw, C.R., and Stone, M.D. (2019). Telomere DNA G-quadruplex folding within actively extending human telomerase. Proc. Natl. Acad. Sci. U. S. A. 116, 9350–9359.

15. Yang, S.Y., Chang, E.Y.C., Lim, J., Kwan, H.H., Monchaud, D., Yip, S., Stirling, P.C., and Wong, J.M.Y. (2021). G-quadruplexes mark alternative lengthening of telomeres. NAR Cancer 3, zcab031.

16. Prorok, P., Artufel, M., Aze, A., Coulombe, P., Peiffer, I., Lacroix, L., Guédin, A., Mergny, J.-L., Damaschke, J., Schepers, A., et al. (2019). Involvement of G-quadruplex regions in mammalian replication origin activity. Nat. Commun. 10, 3274.

17. Mao, S.-Q., Ghanbarian, A.T., Spiegel, J., Martínez Cuesta, S., Beraldi, D., Di Antonio, M., Marsico, G., Hänsel-Hertsch, R., Tannahill, D., and Balasubramanian, S. (2018). DNA G-quadruplex structures mold the DNA methylome. Nat. Struct. Mol. Biol. 25, 951–957.

18. Sarkies, P., Reams, C., Simpson, L.J., and Sale, J.E. (2010). Epigenetic instability due to defective replication of structured DNA. Mol. Cell 40, 703–713.

19. Lopes, J., Piazza, A., Bermejo, R., Kriegsman, B., Colosio, A., Teulade-Fichou, M.-P., Foiani, M., and Nicolas, A. (2011). G-quadruplex-induced instability during leading-strand replication. EMBO J. 30, 4033–4046.

20. Dahan, D., Tsirkas, I., Dovrat, D., Sparks, M.A., Singh, S.P., Galletto, R., and Aharoni, A. (2018). Pif1 is essential for efficient replisome progression through lagging strand G-quadruplex DNA secondary structures. Nucleic Acids Res. 46, 11847–11857.

21. Lerner, L.K., and Sale, J.E. (2019). Replication of G Quadruplex DNA. Genes 10. 10.3390/genes10020095.

22. Belotserkovskii, B.P., Liu, R., Tornaletti, S., Krasilnikova, M.M., Mirkin, S.M., and Hanawalt, P.C. (2010). Mechanisms and implications of transcription blockage by guanine-rich DNA sequences. Proc. Natl. Acad. Sci. U. S. A. 107, 12816–12821.

23. Broxson, C., Beckett, J., and Tornaletti, S. (2011). Transcription arrest by a G quadruplex forming-trinucleotide repeat sequence from the human c-myb gene. Biochemistry 50, 4162–4172.

24. Belotserkovskii, B.P., Neil, A.J., Saleh, S.S., Shin, J.H.S., Mirkin, S.M., and Hanawalt, P.C. (2013). Transcription blockage by homopurine DNA sequences: role of sequence composition and single-strand breaks. Nucleic Acids Res. 41, 1817–1828.

25. Rodriguez, R., Miller, K.M., Forment, J.V., Bradshaw, C.R., Nikan, M., Britton, S., Oelschlaegel, T., Xhemalce, B., Balasubramanian, S., and Jackson, S.P. (2012). Small-molecule-induced DNA damage identifies alternative DNA structures in human genes. Nat. Chem. Biol. 8, 301–310.

26. De Magis, A., Manzo, S.G., Russo, M., Marinello, J., Morigi, R., Sordet, O., and Capranico, G. (2019). DNA damage and genome instability by G-quadruplex ligands are mediated by R loops in human cancer cells. Proc. Natl. Acad. Sci. U. S. A. 116, 816–825.

27. Puget, N., Miller, K.M., and Legube, G. (2019). Non-canonical DNA/RNA structures during Transcription-Coupled Double-Strand Break Repair: Roadblocks or Bona fide repair intermediates? DNA Repair 81, 102661.

28. Georgakopoulos-Soares, I., Morganella, S., Jain, N., Hemberg, M., and Nik-Zainal, S. (2018). Noncanonical secondary structures arising from non-B DNA motifs are determinants of mutagenesis. Genome Res. 28, 1264–1271.

29. Liu, Y., Zhu, X., Wang, K., Zhang, B., and Qiu, S. (2021). The Cellular Functions and Molecular Mechanisms of G-Quadruplex Unwinding Helicases in Humans. Front Mol Biosci 8, 783889.

30. Kunkel, T.A., and Erie, D.A. (2005). DNA mismatch repair. Annu. Rev. Biochem. 74, 681– 710.

31. Larson, E.D., Duquette, M.L., Cummings, W.J., Streiff, R.J., and Maizels, N. (2005). MutSalpha binds to and promotes synapsis of transcriptionally activated immunoglobulin switch regions. Curr. Biol. 15, 470–474.

32. Wu, Y., Shin-ya, K., and Brosh, R.M., Jr (2008). FANCJ helicase defective in Fanconia anemia and breast cancer unwinds G-quadruplex DNA to defend genomic stability. Mol. Cell. Biol. 28, 4116–4128.

33. Sakellariou, D., Bak, S.T., Isik, E., Barroso, S.I., Porro, A., Aguilera, A., Bartek, J., Janscak, P., and Peña-Diaz, J. (2022). MutSβ regulates G4-associated telomeric R-loops to maintain telomere integrity in ALT cancer cells. Cell Rep. 39, 110602.

34. Zeman, M.K., and Cimprich, K.A. (2014). Causes and consequences of replication stress. Nat. Cell Biol. 16, 2–9.

35. Schlacher, K., Christ, N., Siaud, N., Egashira, A., Wu, H., and Jasin, M. (2011). Double-strand break repair-independent role for BRCA2 in blocking stalled replication fork degradation by MRE11. Cell 145, 529–542.

36. Quinet, A., Lemaçon, D., and Vindigni, A. (2017). Replication Fork Reversal: Players and Guardians. Mol. Cell 68, 830–833.

37. Pasero, P., and Vindigni, A. (2017). Nucleases Acting at Stalled Forks: How to Reboot the Replication Program with a Few Shortcuts. Annu. Rev. Genet. 51, 477–499.

38. Rickman, K., and Smogorzewska, A. (2019). Advances in understanding DNA processing and protection at stalled replication forks. J. Cell Biol. 218, 1096–1107.

39. Berti, M., Cortez, D., and Lopes, M. (2020). The plasticity of DNA replication forks in response to clinically relevant genotoxic stress. Nat. Rev. Mol. Cell Biol. 21, 633–651.

40. Sale, J.E. (2013). Translesion DNA synthesis and mutagenesis in eukaryotes. Cold Spring Harb. Perspect. Biol. 5, a012708.

41. Guilliam, T.A., Jozwiakowski, S.K., Ehlinger, A., Barnes, R.P., Rudd, S.G., Bailey, L.J., Skehel, J.M., Eckert, K.A., Chazin, W.J., and Doherty, A.J. (2015). Human PrimPol is a highly error-prone polymerase regulated by single-stranded DNA binding proteins. Nucleic Acids Res. 43, 1056–1068.

42. Ding, H., Descheemaeker, K., Marynen, P., Nelles, L., Carvalho, T., Carmo-Fonseca, M., Collen, D., and Belayew, A. (1996). Characterization of a helicase-like transcription factor involved in the expression of the human plasminogen activator inhibitor-1 gene. DNA Cell Biol. 15, 429–442.

43. Ding, H., Benotmane, A.M., Suske, G., Collen, D., and Belayew, A. (1999). Functional interactions between Sp1 or Sp3 and the helicase-like transcription factor mediate basal expression from the human plasminogen activator inhibitor-1 gene. J. Biol. Chem. 274, 19573–19580.

44. Unk, I., Hajdú, I., Blastyák, A., and Haracska, L. (2010). Role of yeast Rad5 and its human orthologs, HLTF and SHPRH in DNA damage tolerance. DNA Repair 9, 257–267.

45. Unk, I., Hajdú, I., Fátyol, K., Hurwitz, J., Yoon, J.-H., Prakash, L., Prakash, S., and Haracska, L. (2008). Human HLTF functions as a ubiquitin ligase for proliferating cell nuclear antigen polyubiquitination. Proc. Natl. Acad. Sci. U. S. A. 105, 3768–3773.

46. Krijger, P.H.L., Lee, K.-Y., Wit, N., van den Berk, P.C.M., Wu, X., Roest, H.P., Maas, A., Ding, H., Hoeijmakers, J.H.J., Myung, K., et al. (2011). HLTF and SHPRH are not essential for PCNA polyubiquitination, survival and somatic hypermutation: existence of an alternative E3 ligase. DNA Repair 10, 438–444.

47. Lin, J.-R., Zeman, M.K., Chen, J.-Y., Yee, M.-C., and Cimprich, K.A. (2011). SHPRH and HLTF act in a damage-specific manner to coordinate different forms of postreplication repair and prevent mutagenesis. Mol. Cell 42, 237–249.

48. Masuda, Y., Suzuki, M., Kawai, H., Hishiki, A., Hashimoto, H., Masutani, C., Hishida, T., Suzuki, F., and Kamiya, K. (2012). En bloc transfer of polyubiquitin chains to PCNA in vitro is mediated by two different human E2-E3 pairs. Nucleic Acids Res. 40, 10394–10407.

49. Blastyák, A., Hajdú, I., Unk, I., and Haracska, L. (2010). Role of double-stranded DNA translocase activity of human HLTF in replication of damaged DNA. Mol. Cell. Biol. 30, 684–693.

50. Achar, Y.J., Balogh, D., and Haracska, L. (2011). Coordinated protein and DNA remodeling by human HLTF on stalled replication fork. Proc. Natl. Acad. Sci. U. S. A. 108, 14073– 14078.

51. Hishiki, A., Hara, K., Ikegaya, Y., Yokoyama, H., Shimizu, T., Sato, M., and Hashimoto, H. (2015). Structure of a Novel DNA-binding Domain of Helicase-like Transcription Factor (HLTF) and Its Functional Implication in DNA Damage Tolerance. J. Biol. Chem. 290, 13215–13223.

52. Kile, A.C., Chavez, D.A., Bacal, J., Eldirany, S., Korzhnev, D.M., Bezsonova, I., Eichman, B.F., and Cimprich, K.A. (2015). HLTF’s ancient HIRAN domain binds 3’ DNA ends to drive replication fork reversal. Mol. Cell 58, 1090–1100.

53. Chavez, D.A., Greer, B.H., and Eichman, B.F. (2018). The HIRAN domain of helicase-like transcription factor positions the DNA translocase motor to drive efficient DNA fork regression. J. Biol. Chem. 293, 8484–8494.

54. Bai, G., Kermi, C., Stoy, H., Schiltz, C.J., Bacal, J., Zaino, A.M., Hadden, M.K., Eichman, B.F., Lopes, M., and Cimprich, K.A. (2020). HLTF Promotes Fork Reversal, Limiting Replication Stress Resistance and Preventing Multiple Mechanisms of Unrestrained DNA Synthesis. Mol. Cell 78, 1237–1251.e7.

55. Quinet, A., Tirman, S., Jackson, J., Šviković, S., Lemaçon, D., Carvajal-Maldonado, D., González-Acosta, D., Vessoni, A.T., Cybulla, E., Wood, M., et al. (2020). PRIMPOL-Mediated Adaptive Response Suppresses Replication Fork Reversal in BRCA-Deficient Cells. Mol. Cell 77, 461–474.e9.

56. van Toorn, M., Turkyilmaz, Y., Han, S., Zhou, D., Kim, H.-S., Salas-Armenteros, I., Kim, M., Akita, M., Wienholz, F., Raams, A., et al. (2022). Active DNA damage eviction by HLTF stimulates nucleotide excision repair. Mol. Cell 82, 1343–1358.e8.

57. Burkovics, P., Sebesta, M., Balogh, D., Haracska, L., and Krejci, L. (2014). Strand invasion by HLTF as a mechanism for template switch in fork rescue. Nucleic Acids Res. 42, 1711– 1720.

58. Liu, L., Liu, H., Zhou, Y., He, J., Liu, Q., Wang, J., Zeng, M., Yuan, D., Tan, F., Zhou, Y., et al. (2018). HLTF suppresses the migration and invasion of colorectal cancer cells via TGF-β/SMAD signaling in vitro. Int. J. Oncol. 53, 2780–2788.

59. Hamai, Y., Oue, N., Mitani, Y., Nakayama, H., Ito, R., Matsusaki, K., Yoshida, K., Toge, T., and Yasui, W. (2003). DNA hypermethylation and histone hypoacetylation of the HLTF gene are associated with reduced expression in gastric carcinoma. Cancer Sci. 94, 692– 698.

60. Moinova, H.R., Chen, W.-D., Shen, L., Smiraglia, D., Olechnowicz, J., Ravi, L., Kasturi, L., Myeroff, L., Plass, C., Parsons, R., et al. (2002). HLTF gene silencing in human colon cancer. Proc. Natl. Acad. Sci. U. S. A. 99, 4562–4567.

61. Sandhu, S., Wu, X., Nabi, Z., Rastegar, M., Kung, S., Mai, S., and Ding, H. (2012). Loss of HLTF function promotes intestinal carcinogenesis. Mol. Cancer 11, 18.

62. Sirbu, B.M., McDonald, W.H., Dungrawala, H., Badu-Nkansah, A., Kavanaugh, G.M., Chen, Y., Tabb, D.L., and Cortez, D. (2013). Identification of proteins at active, stalled, and collapsed replication forks using isolation of proteins on nascent DNA (iPOND) coupled with mass spectrometry. J. Biol. Chem. 288, 31458–31467.

63. Dungrawala, H., Rose, K.L., Bhat, K.P., Mohni, K.N., Glick, G.G., Couch, F.B., and Cortez, A. (2015). The Replication Checkpoint Prevents Two Types of Fork Collapse without Regulating Replisome Stability. Mol. Cell 59, 998–1010.

64. Duquette, M.L., Handa, P., Vincent, J.A., Taylor, A.F., and Maizels, N. (2004). Intracellular transcription of G-rich DNAs induces formation of G-loops, novel structures containing G4 DNA. Genes Dev. 18, 1618–1629.

65. Crossley, M.P., Bocek, M., and Cimprich, K.A. (2019). R-Loops as Cellular Regulators and Genomic Threats. Mol. Cell 73, 398–411.

66. Petermann, E., Lan, L., and Zou, L. (2022). Sources, resolution and physiological relevance of R-loops and RNA-DNA hybrids. Nat. Rev. Mol. Cell Biol. 23, 521–540.

67. Owen, B.A.L., Yang, Z., Lai, M., Gajec, M., Badger, J.D., 2nd, Hayes, J.J., Edelmann, W., Kucherlapati, R., Wilson, T.M., and McMurray, C.T. (2005). (CAG)(n)-hairpin DNA binds to Msh2-Msh3 and changes properties of mismatch recognition. Nat. Struct. Mol. Biol. 12, 663–670.

68. Young, S.J., Sebald, M., Shah Punatar, R., Larin, M., Masino, L., Rodrigo-Brenni, M.C., Liang, C.-C., and West, S.C. (2020). MutSβ Stimulates Holliday Junction Resolution by the SMX Complex. Cell Rep. 33, 108289.

69. Burdova, K., Mihaljevic, B., Sturzenegger, A., Chappidi, N., and Janscak, P. (2015). The Mismatch-Binding Factor MutSβ Can Mediate ATR Activation in Response to DNA Double-Strand Breaks. Mol. Cell 59, 603–614.

70. McKinney, J.A., Wang, G., Mukherjee, A., Christensen, L., Subramanian, S.H.S., Zhao, J., and Vasquez, K.M. (2020). Distinct DNA repair pathways cause genomic instability at alternative DNA structures. Nat. Commun. 11, 236.

71. Mengoli, V., Ceppi, I., Sanchez, A., Cannavo, E., Halder, S., Scaglione, S., Gaillard, P.-H., McHugh, P.J., Riesen, N., Pettazzoni, P., et al. (2023). WRN helicase and mismatch repair complexes independently and synergistically disrupt cruciform DNA structures. EMBO J. 42, e111998.

72. Orlando, D.A., Chen, M.W., Brown, V.E., Solanki, S., Choi, Y.J., Olson, E.R., Fritz, C.C., Bradner, J.E., and Guenther, M.G. (2014). Quantitative ChIP-Seq normalization reveals global modulation of the epigenome. Cell Rep. 9, 1163–1170.

73. Hui, W.W.I., Simeone, A., Zyner, K.G., Tannahill, D., and Balasubramanian, S. (2021). Single-cell mapping of DNA G-quadruplex structures in human cancer cells. Sci. Rep. 11, 23641.

74. Marsico, G., Chambers, V.S., Sahakyan, A.B., McCauley, P., Boutell, J.M., Antonio, M.D., and Balasubramanian, S. (2019). Whole genome experimental maps of DNA G-quadruplexes in multiple species. Nucleic Acids Res. 47, 3862–3874.

75. Tan, J., Wang, X., Phoon, L., Yang, H., and Lan, L. (2020). Resolution of ROS-induced G-quadruplexes and R-loops at transcriptionally active sites is dependent on BLM helicase. FEBS Lett. 594, 1359–1367.

76. Lim, G., and Hohng, S. (2020). Single-molecule fluorescence studies on cotranscriptional G-quadruplex formation coupled with R-loop formation. Nucleic Acids Res. 48, 9195–9203.

77. Crossley, M.P., Brickner, J.R., Song, C., Zar, S.M.T., Maw, S.S., Chédin, F., Tsai, M.-S., and Cimprich, K.A. (2021). Catalytically inactive, purified RNase H1: A specific and sensitive probe for RNA-DNA hybrid imaging. J. Cell Biol. 220. 10.1083/jcb.202101092.

78. Garcia-Barcena, C., Osinalde, N., Ramirez, J., and Mayor, U. (2020). How to Inactivate Human Ubiquitin E3 Ligases by Mutation. Front Cell Dev Biol 8, 39.

79. Yusufzai, T., and Kadonaga, J.T. (2008). HARP is an ATP-driven annealing helicase. Science 322, 748–750.

80. Arora, R., Lee, Y., Wischnewski, H., Brun, C.M., Schwarz, T., and Azzalin, C.M. (2014). RNaseH1 regulates TERRA-telomeric DNA hybrids and telomere maintenance in ALT tumour cells. Nat. Commun. 5, 5220.

81. Santos-Pereira, J.M., and Aguilera, A. (2015). R loops: new modulators of genome dynamics and function. Nat. Rev. Genet. 16, 583–597.

82. García-Muse, T., and Aguilera, A. (2019). R Loops: From Physiological to Pathological Roles. Cell 179, 604–618.

83. Amato, R., Valenzuela, M., Berardinelli, F., Salvati, E., Maresca, C., Leone, S., Antoccia, A., and Sgura, A. (2020). G-quadruplex Stabilization Fuels the ALT Pathway in ALT-positive Osteosarcoma Cells. Genes 11. 10.3390/genes11030304.

84. Barroso-González, J., García-Expósito, L., Galaviz, P., Lynskey, M.L., Allen, J.A.M., Hoang, S., Watkins, S.C., Pickett, H.A., and O’Sullivan, R.J. (2021). Anti-recombination function of MutSα restricts telomere extension by ALT-associated homology-directed repair. Cell Rep. 37, 110088.

85. Lezaja, A., Panagopoulos, A., Wen, Y., Carvalho, E., Imhof, R., and Altmeyer, M. (2021). RPA shields inherited DNA lesions for post-mitotic DNA synthesis. Nat. Commun. 12, 3827.

86. Zhang, J.-M., and Zou, L. (2020). Alternative lengthening of telomeres: from molecular mechanisms to therapeutic outlooks. Cell Biosci. 10, 30.

87. Loe, T.K., Li, J.S.Z., Zhang, Y., Azeroglu, B., Boddy, M.N., and Denchi, E.L. (2020). Telomere length heterogeneity in ALT cells is maintained by PML-dependent localization of the BTR complex to telomeres. Genes Dev. 34, 650–662.

88. Kumar, C., Batra, S., Griffith, J.D., and Remus, D. (2021). The interplay of RNA:DNA hybrid structure and G-quadruplexes determines the outcome of R-loop-replisome collisions. Elife 10. 10.7554/eLife.72286.

89. Williams, S.L., Casas-Delucchi, C.S., Raguseo, F., Guneri, D., Li, Y., Minamino, M., Fletcher, E.E., Yeeles, J.T.P., Keyser, U.F., Waller, Z.A.E., et al. (2023). Replication-induced DNA secondary structures drive fork uncoupling and breakage. bioRxiv, 2022.11.18.517070. 10.1101/2022.11.18.517070.

90. Schiavone, D., Jozwiakowski, S.K., Romanello, M., Guilbaud, G., Guilliam, T.A., Bailey, L.J., Sale, J.E., and Doherty, A.J. (2016). PrimPol Is Required for Replicative Tolerance of G Quadruplexes in Vertebrate Cells. Mol. Cell 61, 161–169.

91. Gan, W., Guan, Z., Liu, J., Gui, T., Shen, K., Manley, J.L., and Li, X. (2011). R-loop-mediated genomic instability is caused by impairment of replication fork progression. Genes Dev. 25, 2041–2056.

92. Xu, H., Di Antonio, M., McKinney, S., Mathew, V., Ho, B., O’Neil, N.J., Santos, N.D., Silvester, J., Wei, V., Garcia, J., et al. (2017). CX-5461 is a DNA G-quadruplex stabilizer with selective lethality in BRCA1/2 deficient tumours. Nat. Commun. 8, 14432.

93. Hodson, C., van Twest, S., Dylewska, M., O’Rourke, J.J., Tan, W., Murphy, V.J., Walia, M., Abbouche, L., Nieminuszczy, J., Dunn, E., et al. (2022). Branchpoint translocation by fork remodelers as a general mechanism of R-loop removal. Cell Rep. 41, 111749.

94. Déjardin, J., and Kingston, R.E. (2009). Purification of proteins associated with specific genomic Loci. Cell 136, 175–186.

95. Lee, W.T.C., Yin, Y., Morten, M.J., Tonzi, P., Gwo, P.P., Odermatt, D.C., Modesti, M., Cantor, S.B., Gari, K., Huang, T.T., et al. (2021). Single-molecule imaging reveals replication fork coupled formation of G-quadruplex structures hinders local replication stress signaling. Nat. Commun. 12, 2525.

96. Frizzell, A., Nguyen, J.H.G., Petalcorin, M.I.R., Turner, K.D., Boulton, S.J., Freudenreich, C.H., and Lahue, R.S. (2016). RTEL1 Inhibits Trinucleotide Repeat Expansions and Fragility. Cell Rep. 16, 2047.

97. Rastokina, A., Cebrián, J., Mozafari, N., Mandel, N.H., Smith, C.I.E., Lopes, M., Zain, R., and Mirkin, S.M. (2023). Large-scale expansions of Friedreich’s ataxia GAA•TTC repeats in an experimental human system: role of DNA replication and prevention by LNA-DNA oligonucleotides and PNA oligomers. Nucleic Acids Res. 10.1093/nar/gkad441.

98. Yang, Q., Zhang, R., Wang, X.W., Linke, S.P., Sengupta, S., Hickson, I.D., Pedrazzi, G., Perrera, C., Stagljar, I., Littman, S.J., et al. (2004). The mismatch DNA repair heterodimer, hMSH2/6, regulates BLM helicase. Oncogene 23, 3749–3756.

99. Saydam, N., Kanagaraj, R., Dietschy, T., Garcia, P.L., Peña-Diaz, J., Shevelev, I., Stagljar, I., and Janscak, P. (2007). Physical and functional interactions between Werner syndrome helicase and mismatch-repair initiation factors. Nucleic Acids Res. 35, 5706–5716.

100. Peng, M., Litman, R., Xie, J., Sharma, S., Brosh, R.M., Jr, and Cantor, S.B. (2007). The FANCJ/MutLalpha interaction is required for correction of the cross-link response in FA-J cells. EMBO J. 26, 3238–3249.

101. Peng, M., Xie, J., Ucher, A., Stavnezer, J., and Cantor, S.B. (2014). Crosstalk between BRCA-Fanconi anemia and mismatch repair pathways prevents MSH2-dependent aberrant DNA damage responses. EMBO J. 33, 1698–1712.

102. Peng, M., Cong, K., Panzarino, N.J., Nayak, S., Calvo, J., Deng, B., Zhu, L.J., Morocz, M., Hegedus, L., Haracska, L., et al. (2018). Opposing Roles of FANCJ and HLTF Protect Forks and Restrain Replication during Stress. Cell Rep. 24, 3251–3261.

103. Gray, L.T., Vallur, A.C., Eddy, J., and Maizels, N. (2014). G quadruplexes are genomewide targets of transcriptional helicases XPB and XPD. Nat. Chem. Biol. 10, 313–318.

104. Wessel, S.R., Mohni, K.N., Luzwick, J.W., Dungrawala, H., and Cortez, D. (2019). Functional Analysis of the Replication Fork Proteome Identifies BET Proteins as PCNA Regulators. Cell Rep. 28, 3497–3509.e4.

105. Tyanova, S., Temu, T., Sinitcyn, P., Carlson, A., Hein, M.Y., Geiger, T., Mann, M., and Cox, J. (2016). The Perseus computational platform for comprehensive analysis of (prote)omics data. Nat. Methods 13, 731–740.

106. Spencley, A.L., Bar, S., Swigut, T., Flynn, R.A., Lee, C.H., Chen, L.-F., Bassik, M.C., and Wysocka, J. (2023). Co-transcriptional genome surveillance by HUSH is coupled to termination machinery. Mol. Cell 83, 1623–1639.e8.

107. Langmead, B., and Salzberg, S.L. (2012). Fast gapped-read alignment with Bowtie 2. Nat. Methods 9, 357–359.

108. Amemiya, H.M., Kundaje, A., and Boyle, A.P. (2019). The ENCODE Blacklist: Identification of Problematic Regions of the Genome. Sci. Rep. 9, 9354.

109. Ramírez, F., Ryan, D.P., Grüning, B., Bhardwaj, V., Kilpert, F., Richter, A.S., Heyne, S., Dündar, F., and Manke, T. (2016). deepTools2: a next generation web server for deep-sequencing data analysis. Nucleic Acids Res. 44, W160–5.

110. Zhang, Y., Liu, T., Meyer, C.A., Eeckhoute, J., Johnson, D.S., Bernstein, B.E., Nusbaum, C., Myers, R.M., Brown, M., Li, W., et al. (2008). Model-based analysis of ChIP-Seq (MACS). Genome Biol. 9, R137.

111. Landt, S.G., Marinov, G.K., Kundaje, A., Kheradpour, P., Pauli, F., Batzoglou, S., Bernstein, B.E., Bickel, P., Brown, J.B., Cayting, P., et al. (2012). ChIP-seq guidelines and practices of the ENCODE and modENCODE consortia. Genome Res. 22, 1813–1831.

112. Quinlan, A.R. (2014). BEDTools: The Swiss-Army Tool for Genome Feature Analysis. Curr. Protoc. Bioinformatics 47, 11.12.1-34.

113. Heinz, S., Benner, C., Spann, N., Bertolino, E., Lin, Y.C., Laslo, P., Cheng, J.X., Murre, C., Singh, H., and Glass, C.K. (2010). Simple combinations of lineage-determining transcription factors prime cis-regulatory elements required for macrophage and B cell identities. Mol. Cell 38, 576–589.

114. Campbell, S.J., Edwards, R.A., Leung, C.C.Y., Neculai, D., Hodge, C.D., Dhe-Paganon, S., and Glover, J.N.M. (2012). Molecular insights into the function of RING finger (RNF)-containing proteins hRNF8 and hRNF168 in Ubc13/Mms2-dependent ubiquitylation. J. Biol. Chem. 287, 23900–23910.

115. Kiianitsa, K., Solinger, J.A., and Heyer, W.-D. (2003). NADH-coupled microplate photometric assay for kinetic studies of ATP-hydrolyzing enzymes with low and high specific activities. Anal. Biochem. 321, 266–271.

